# Phase transition of bacterial single-stranded DNA binding (SSB) protein upon stress response and metabolic adaptation

**DOI:** 10.1101/2025.08.28.672810

**Authors:** Péter Ecsédi, Júlia Szittyai, János Pálinkás, Bálint Jezsó, Viktoria Katran, Zoltán J. Kovács, Henriett Halász, Tasvilla Sonallya, Tünde Juhász, Tamás Beke-Somfai, Szilvia Barkó, Edina Szabó-Meleg, Mihály Kovács

## Abstract

Single-stranded DNA-binding proteins (SSBs) are ubiquitous factors of genome metabolism, recently recognized for their ability to undergo liquid–liquid phase separation (LLPS). While *Escherichia coli* SSB (EcSSB) has emerged as a model for bacterial LLPS *in vitro*, its phase behavior and functional dynamics *in vivo* have remained largely unexplored. Here, we define the subcellular organization of EcSSB under diverse physiological and stress conditions using super-resolution microscopy coupled with newly developed image analysis tools. We show that EcSSB forms dynamic intracellular assemblies during exponential growth, which partially dissolve in response to DNA damage, oxidative stress, antibiotic exposure, and metabolic adaptation. In contrast, these dynamic reorganizations are attenuated in stationary phase cells. Moreover, we show that EcSSB foci exhibit limited overlap with nucleoid regions under stress-free conditions, whereas stress induction is accompanied by increased DNA association. These *in vivo* observations are consistent with stress- and growth-phase dependent modulation of EcSSB organization and together with biophysical characterization of EcSSB condensation *in vitro* suggest a role for LLPS-based dynamic storage and mobilization in response to physiological demands. Our work provides a quantitative framework for analyzing the cellular organization of bacterial proteins and the spatial regulation of genome maintenance factors in changing environments. This knowledge may also support future strategies targeting SSB organization pathways for antimicrobial development.

## Introduction

Living cells must respond rapidly and effectively to DNA damage to mitigate its potentially severe impacts. Among others, ionizing and ultraviolet radiation, alkylating agents, and polycyclic aromatic hydrocarbons can cause exogenous DNA damage ^1^. In addition, DNA is also vulnerable during physiological DNA metabolic processes including DNA replication, recombination, and repair. Nearly all genome maintenance pathways involve exposure of single-stranded DNA (ssDNA) segments whose processing is mediated by the action of single-stranded DNA binding (SSB) proteins in both prokaryotic and eukaryotic cells ^2^.

SSBs protect ssDNA from degradation and prevent its reannealing. In addition, SSBs recruit and organize genome maintenance complexes through protein–protein interactions (PPIs) ^2^. *Escherichia coli* SSB (EcSSB), the prototypic SSB, is a homotetramer of 178-amino acid (19-kDa) subunits ^3^. An EcSSB subunit comprises the N-terminal globular oligonucleotide/oligosaccharide-binding (OB) domain (amino acids (aa) 1-113) responsible for ssDNA binding ^4^, and an intrinsically disordered linker (IDL, aa 114-178) driving PPIs ^5,6^. The C-terminal segment of the IDL (so-called C-terminal peptide, CTP, aa 170-178), a key PPI element, is highly conserved among bacteria ^2,3,5,7,8^. It was recently demonstrated that EcSSB liquid-liquid phase separation (LLPS) is brought about by a fine-tuned interplay of the protein’s different structural regions ^9–11^.

LLPS of proteins has been known for more than a century, yet its biological importance has only been recently recognized ^12–15^. Numerous proteins have been reported to be capable of LLPS-based condensation ^12,16^, and LLPS has emerged as a key mechanism for the formation of membraneless organelles ^17,18^. Examples for these structures in eukaryotes include Cajal bodies, P granules, and the nucleolus ^15^. The formation of LLPS particles is typically driven by weak multivalent protein-protein and/or protein-nucleic acid interactions ^19,20^. LLPS condensates serve a wide range of functions, such as buffering concentrations of proteins, rapid mobilization of biomolecules upon stress, and enhancing reactions by selectively concentrating molecules involved in biochemical processes ^21,22^. Although the role of protein condensation in bacteria has remained less explored, it has recently become increasingly apparent that this phenomenon bears special importance for bacteria lacking internal membrane-based compartmentalization ^12,21^.

The above-mentioned recognition aligns with our previous *in vitro* finding that increasing concentrations of ssDNA dissolve EcSSB condensates in a stoichiometric fashion, suggesting a role for dynamic EcSSB decondensation in bacterial genome maintenance ^9^. Intracellular EcSSB assemblies have been generally attributed to replication-associated complexes ^6,23–28^. However, various observations suggest that replication association alone may not fully explain all aspects of EcSSB organization. A substantial fraction of EcSSB assemblies do not colocalize with replisome components ^24,26^ or with nucleoids ^6^. Furthermore, EcSSB foci have been reported to undergo fusion-like events and changes in intensity over time ^26^, behaviors that are consistent with, although not exclusive to, condensation-driven organization. In addition, Reyes-Lamothe et al. reported that cells frequently contained a single prominent EcSSB focus prior to replication initiation ^23^, a finding that is difficult to reconcile solely with replication-associated EcSSB. A main challenge in studying intracellular protein assemblies in bacteria is the difficulty of visualization at this scale. Using super-resolution microscopy, Zhao *et al.* achieved highly precise intracellular localization of EcSSB in an *E. coli* strain expressing a GFP-tagged EcSSB variant from its native chromosomal locus ^29^. In the unstressed state, EcSSB foci localized near the membrane and, to a lesser extent, at genomic sites. Treatment with the DNA damaging agent mitomycin C triggered the dissociation of EcSSB from the membrane, with nearly all EcSSB being found inside the cell.

The above-mentioned observations suggest a sequestration mechanism that, in light of more recent *in vitro* evidence, could be mediated by LLPS. Taken together, these findings, along with the ssDNA-regulated condensation features of EcSSB observed *in vitro* ^9–11,30^, motivated us to perform a comprehensive, super-resolution scale definition of EcSSB subcellular patterns during bacterial growth, stress responses, and metabolic adaptations, taking into account its LLPS propensity. Therefore, in this work we combine super-resolution microscopy and quantitative image analysis to characterize stress- and growth-phase dependent remodeling of EcSSB in cells, correlating with DNA damage, metabolic state, and proliferation cues. Our results are consistent with a model involving dynamic SSB condensation that regulates protein organization, macromolecular mobility, sequestration, and release in bacteria.

## Results

### Characterization of the EcSSB-GFP expressing cell line

In this work we used an *E. coli* strain #29003 (MG1655, ΔlacY::kan flgE:EcSSB-GFP-cam), which was a gift from Piero R. Bianco ^29^. Importantly, the sequence encoding EcSSB-GFP was introduced downstream—not in place—of the wild-type (WT) EcSSB coding sequence. Consequently, these bacteria express both WT EcSSB and EcSSB-GFP without induction, as established by Western blot analysis ^29^. Previously it was shown that, under these conditions, both WT and chimeric SSB tetramers can form, with the latter retaining WT function even when the tetramer contains up to two GFP-fused subunits ^31^. Thus, the fused construct can be utilized for fluorescence-based detection, while the WT protein provides intact SSB functionality in cells ^29^.

To further assess the effects of EcSSB-GFP expression, we compared the growth and morphology of these cells with those of a control cell line (#27869: MG1655 ΔlacY::kan), which is otherwise identical to #29003 but does not express EcSSB-GFP ^29^ (**Fig. S1**). We found no detectable differences in cell morphology or growth between these cell lines under normal conditions. Upon exposure to typical stressors, both cell lines exhibited highly similar growth profiles (**Fig. S1C**). Notably, cells expressing EcSSB-GFP alongside the WT protein showed slightly smaller extents of stress-induced growth suppression, presumably due to the additional EcSSB-GFP expression.

Notably, cells expressing EcSSB C-terminal fluorescent protein (CFP, YPet) fusions, similar to the one used in the current study, have also shown unperturbed growth and cell cycle parameters ^23,25^. Moreover, independent studies employing alternative fluorescent SSB fusion designs (mTur2 or GFP inserted within the IDL region) have reported highly similar intracellular localization patterns ^6,26,28^. In addition, fluorescence recovery after photobleaching (FRAP) experiments revealed rapid molecular exchange within EcSSB-YPet assemblies, indicating that these structures are dynamic rather than static protein aggregates ^25^. Collectively, these observations indicate that intracellular EcSSB assemblies are reproducibly observed across distinct labeling strategies and are unlikely to arise from fusion-specific artifacts.

### EcSSB forms dynamic assemblies during growth, which consolidate upon growth arrest

To characterize the subcellular distribution of EcSSB during bacterial growth, we recorded subcellular EcSSB patterns in exponential, stationary, and late (“death”) growth phases (**Fig. 1 and Table S1**). In the early exponential phase (*OD*_600_ = 0.2), cells exhibited EcSSB-GFP signals with only a few faint foci. The majority of EcSSB assemblies (typically 1, but up to 6, per cell) were observed in the mid exponential growth phase (*OD*_600_ = 0.6). Notably, in late exponential and early stationary phases (*OD*_600_ = 0.6 + 1.5 h, + 3 h, and + 4.5 h), the number of EcSSB foci in cells tended to consolidate into a single dominant focus, with cells lacking detectable foci becoming exceedingly rare. This trend shifted in later phases (*OD*_600_ = 0.6 + 7 h, + 10 h, and + 24 h), where the dominant EcSSB assemblies progressively diminished. During these stages, the overall EcSSB-GFP signal also decreased, consistent with entry into long-term stationary phase and cell death ^32,33^.

**Figure 1.**
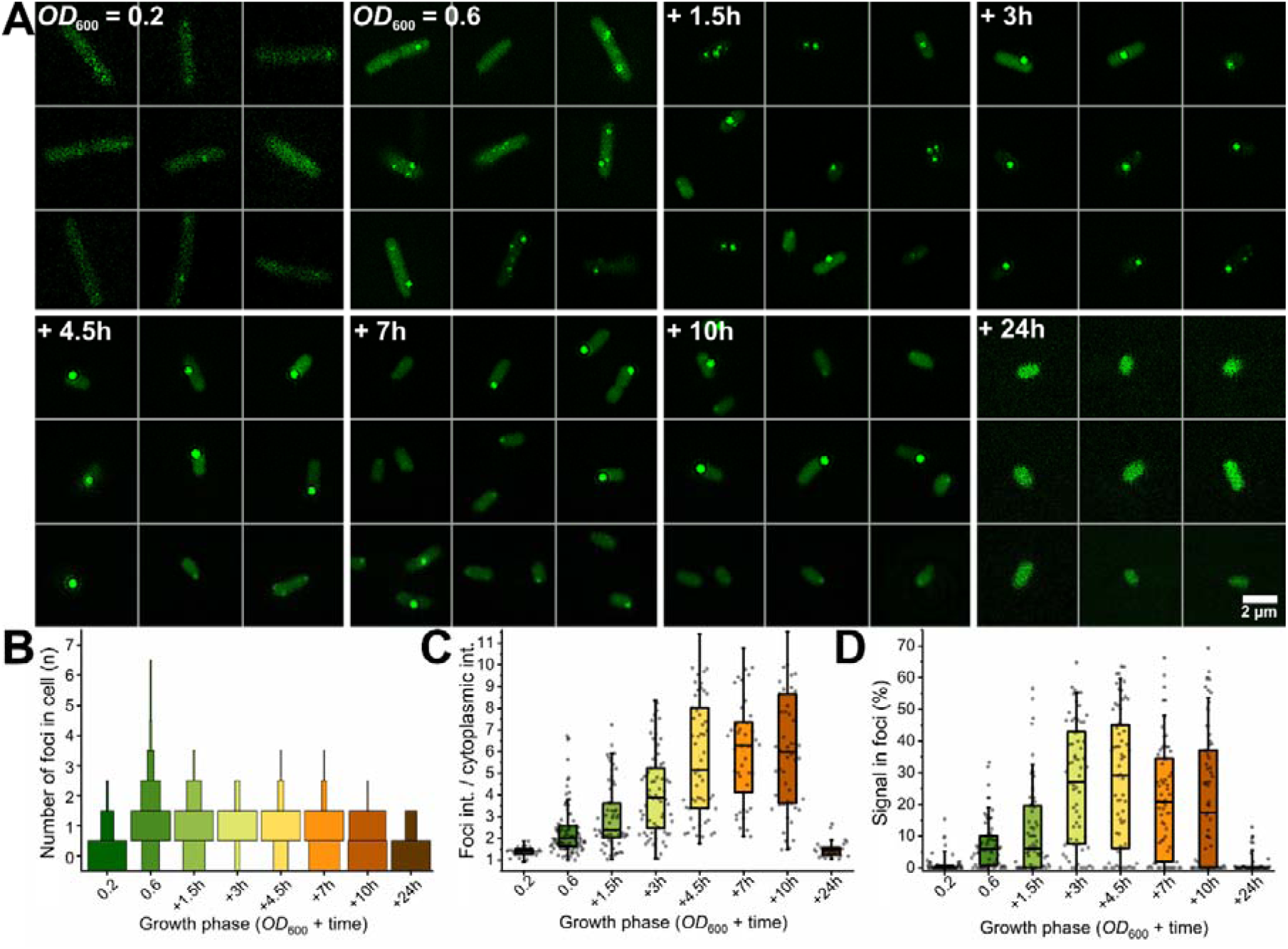
EcSSB shows growth-phase dependent intracellular organization. **(A)** Structured illumination microscopy (SIM) images of the EcSSB-GFP expressing E. coli strain #29003 ^29^ in different growth phases. Images were evaluated using a newly developed analysis method described in **Supplementary file 1**. Quantitative analysis of **(B)** the number of foci per cell (bin size = 1), **(C)** their intensities, and **(D)** the percentage of signal contained in discrete foci reveals growth-phase dependent changes in EcSSB organization, characterized by a gradual reduction in the number of detectable fluorescent foci and an increase in signal intensity in remaining structures (n = 65 cells analyzed per condition, see also **Table S1**).

The time-dependent trend in EcSSB-GFP foci signal intensity indicates that the appearance of more intense foci is associated with slower growth conditions (*e.g. OD*_600_ = 0.6 compared to *OD*_600_ = 0.6 + 3 h, **Fig. 1**). Furthermore, as the number of discrete signals decreased, their intensity increased, reflecting a redistribution of EcSSB into fewer but more pronounced intracellular assemblies. This observation is consistent with a shift in the spatial organization of EcSSB over time. Notably, similar trends are observed *in vitro* during EcSSB condensation behavior under comparable conditions (see *in vitro* biophysical data described below). Based on EcSSB-GFP signal intensity analysis including all cells either harboring or lacking fluorescence foci, we detected a reduction in diffuse cytoplasmic EcSSB-GFP signal as cultures grew older, further indicating the assimilation of the dissolved EcSSB pool (eventually, up to 70 % of the total EcSSB-GFP signal became localized in foci) (**Fig. 1** and **Fig. S2**). We note that the high proportion of the cellular EcSSB-GFP signal contained in foci is also indicative of efficient incorporation of EcSSB-GFP into endogenous (wild-type) EcSSB assemblies (see also below for *in vitro* enrichment of EcSSB-GFP in WT EcSSB LLPS condensates). During the death phase (*OD*_600_ = 0.6 + 10 h and 24 h), the overall cellular EcSSB-GFP signal declined, ultimately leading to the complete disappearance of discrete signals.

### EcSSB-GFP assemblies disassemble under stress, but this process is attenuated in stationary-phase cells

Next, we analyzed the effects of a broad panel of stressors and metabolic modulators (described in **Table S2**, including dosages and mechanisms of action) on cells in the exponential growth phase (*OD*_600_ = 0.6). Of the applied agents, (*i*) hydroxyurea (HU) and NaCl suppress, while (*ii*) L-arginine (L-Arg), glucose, and lactose stimulate cell growth; (*iii*) UV irradiation, nitrofurantoin, and methyl methanesulfonate (MMS) directly damage DNA; (*iv*) diamide and H_2_O_2_ cause different forms of oxidative stress; (*v*) myricetin and taxifolin are antibacterial flavonoids interacting with prokaryotic SSBs ^31,32^ and (*vi*) MES (2-(N-morpholino)ethanesulfonic acid, pH 6) was used to cause pH stress. Importantly, most applied stressors induced a decrease in the number and intensity of EcSSB-GFP foci, with a concomitant increase in cytoplasmic EcSSB-GFP signal (**Fig. 2 and Table S3**). UV, L-Arg, glucose, H_2_O_2_, MES pH 6, myricetin, and taxifolin exerted the strongest effects. Upon these treatments, the most frequently observed number of foci dropped to 0 (except in case of H_2_O_2_). Moreover, with a few exceptions, stressed cells contained less than 5 % of the EcSSB-GFP signal (if any) in discrete signals, in contrast to the typical 10-20 % (maximum 40 %) in untreated cells (**Fig. 2D**). Notably, glucose, which is a more favored energy source for *E. coli* than lactose, exerted stronger reduction in EcSSB-GFP foci compared to the sugar alternative (**Fig. 2 and Table S3**). Other stressors including HU, MMS and nitrofurantoin caused mild or no changes in EcSSB-GFP foci, while NaCl was associated with an increase in EcSSB-GFP foci (**Fig. 2 and Table S3**). These results indicate stress- and metabolism-dependent changes in EcSSB organization. Diamide had a special effect in causing EcSSB assemblies to localize more toward the cell membrane, similar to that seen for untreated cells by Zhao *et al.* ^29^, in line with the observed interaction of EcSSB with membrane phospholipids in that study.

**Figure 2.**
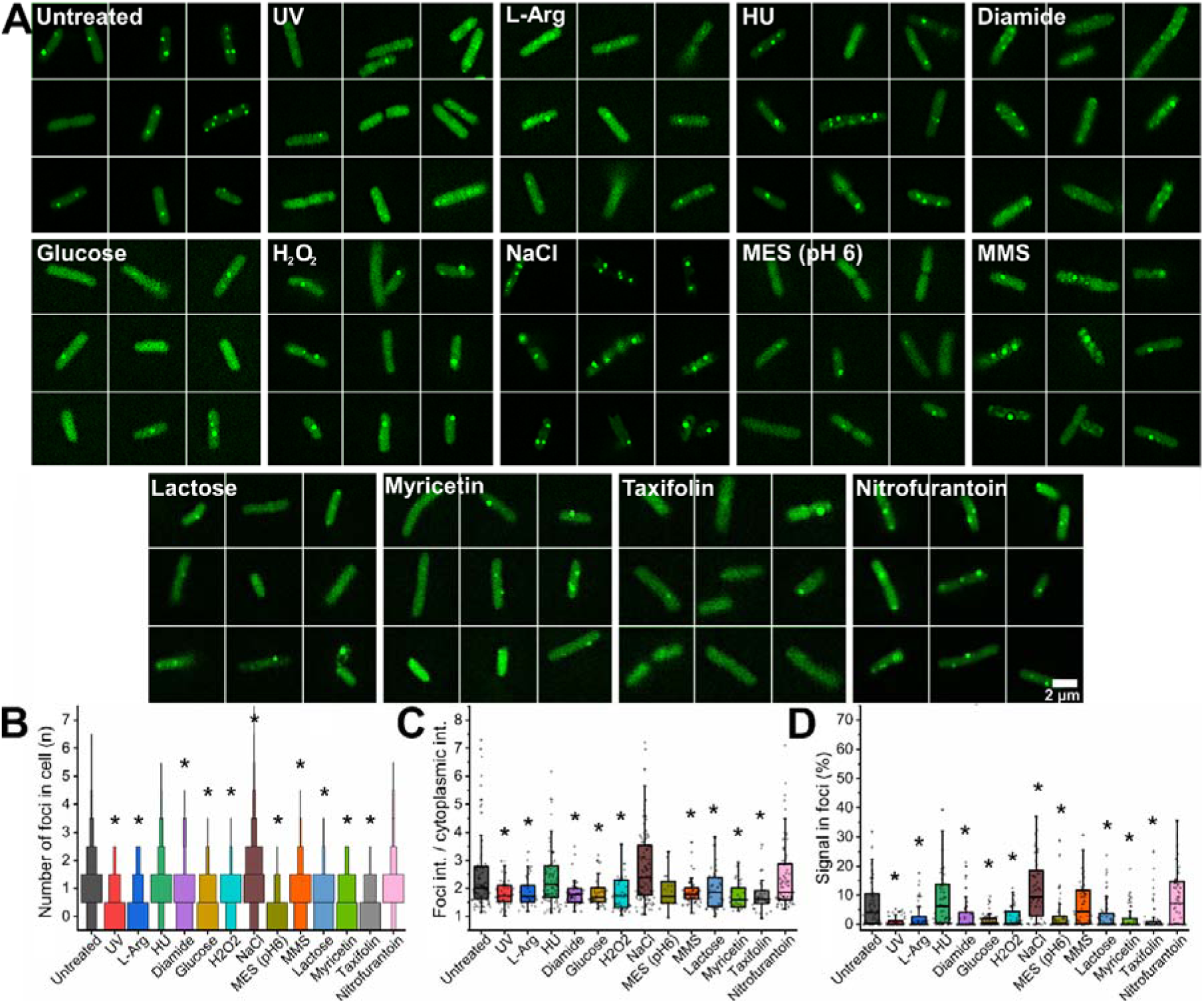
Various stressors induce a reduction of EcSSB-GFP foci in exponential growth phase (OD_600_ = 0.6). **(A)** Representative SIM images of E. coli strain #29003 show distinct EcSSB-GFP foci in untreated cells. Upon exposure to various stressors (as indicated; see **Table S2** for conditions), the EcSSB-GFP signal became dispersed throughout the cytoplasm, while **(B)** the number (bin size = 1) and **(C)** intensity of foci decreased. **(D)** The reduction in the percentage of signal in foci further corroborates stress-induced reorganization of EcSSB. Asterisks denote significant difference compared to untreated cells (Mann-Whitney test, p < 0.05, n = 65 cells analyzed per condition, see also **Table S3**).

Here we note that we also tested the effect of the FM 4-64 membrane dye, used by Zhao *et al*. ^29^, on EcSSB-GFP subcellular patterns. Intriguingly, we found that the dye caused EcSSB-GFP localization to membrane-adjacent cellular areas, suggesting a previously unrecognized nonspecific effect (**Figs. S3 E-G**), and therefore we refrained from its use in other experiments. The possibility that the membrane-associated green signal could originate from FM 4-64 fluorescence bleed-through can be excluded based on (*i*) the complete disappearance of EcSSB-GFP signal from internal cellular areas; (*ii*) the lack of the EcSSB-GFP membrane localization effect at lower FM 4-64 concentrations (2 µM, as used by Zhao *et al.* ^29^) with detectable red fluorescence; (*iii*) the occasional presence of red fluorescent spots (FM 4-64 aggregates) showing no signal in the green channel.

Next, we determined the effects of the above-mentioned stressors during the stationary and death phases (*OD*_600_ = 0.6 + 3h (**Fig. 3 and Table S4**), *OD*_600_ = 0.6 + 24 h (**Fig. S3 and Table S5**), respectively). In the stationary phase, stressors elicited generally milder responses compared to the exponential phase. Nevertheless, UV, L-Arg, glucose, H_2_O_2_, and MES (pH 6) still elicited effective, albeit partial changes in EcSSB-GFP organization. These changes were characterized by an increase in diffuse cytoplasmic EcSSB-GFP signal and a redistribution of signal into more numerous but less intense foci. Glucose, lactose and, notably, HU were exceptions, as these agents primarily increased the cytoplasmic signal with limited changes in the number of detectable foci.

**Figure 3.**
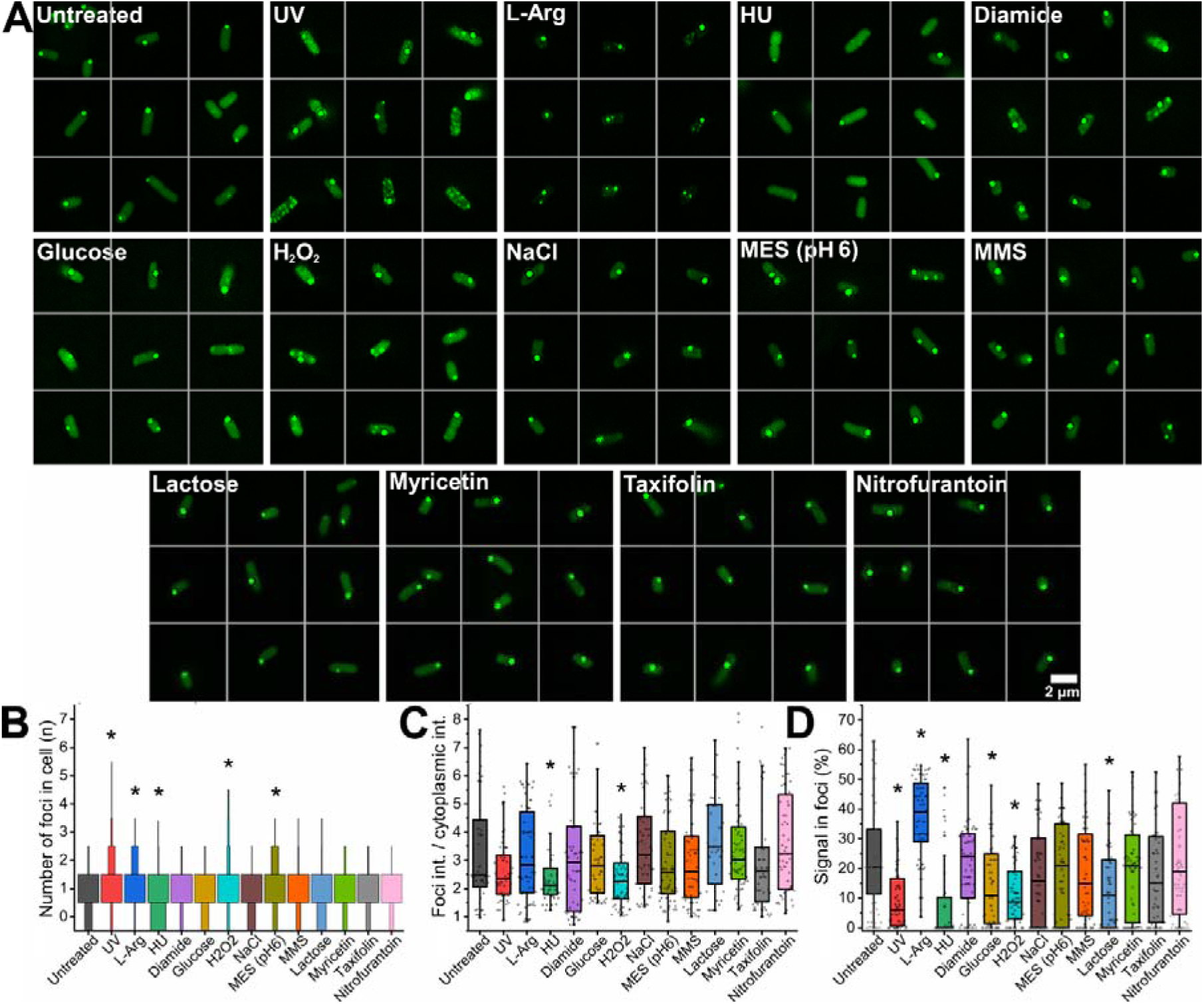
Stress-induced changes in EcSSB-GFP spatial organization are attenuated in stationary growth phase (OD_600_ = 0.6 + 3 h). (A) Representative SIM images of E. coli strain #29003 show a single bright focus in each cell under stationary-phase conditions. Stressors affected EcSSB-GFP patterns to a markedly lesser extent than in exponential-phase cells (OD_600_ = 0.6, **Fig. 2**; see Table S2 for conditions). However, UV, MES (pH 6), H_2_O_2_, and glucose remained the most effective disruptors of EcSSB assemblies. Analysis of (B) the number (bin size = 1) and (C) intensity of foci in cells showed that stressors causing DNA damage (e.g. UV and H_2_O_2_) elicit a rapid response initiating the fragmentation of EcSSB assemblies, while glucose gradually decreases foci intensity. (D) Cells contained markedly higher percentages of EcSSB-GFP signal in discrete foci compared to cells in earlier phases (OD_600_ = 0.2 and 0.6, **Figs. 1-2**). Asterisks denote significant difference compared to untreated cells (Mann-Whitney test, p < 0.05, n = 65 cells analyzed per condition, see also Table S4).

Unlike exponential-phase cells, stationary-phase cells retained up to 70 % of their EcSSB-GFP signal in discrete foci. However, in the presence of stressors including glucose, H_2_O_2_, HU, and UV, this fraction often dropped below 40 %. In addition, diamide exhibited a reduced propensity to induce redistribution of signal toward membrane-associated regions compared to that seen in the exponential phase, while UV occasionally promoted such localization (**Figs. 2 and 3**). The latter effect was not detectable in exponential-phase experiments, as UV treatment led to the strong reduction of detectable EcSSB-GFP foci in that phase (**Fig. 2**). Nonetheless, this UV-induced behavior was less pronounced in the stationary phase compared to diamide during the exponential phase. Overall, stationary-phase cells showed altered stress-dependent changes in EcSSB-GFP spatial distribution compared to exponential-phase cells.

The subcellular localization of EcSSB-GFP underwent significant changes in the death phase (**Fig. S3 and Table S5**), with only faint cytoplasmic signal and few or no detectable foci. This phenomenon may occur because cells begin to reutilize their resources, while also degrading their EcSSB pool. Most stressors, with the exception of NaCl, diamide, and nitrofurantoin, had minimal effects on EcSSB-GFP localization in this phase. In contrast, these conditions were associated with the presence of small, faint foci.

### Stress induction increases spatial association between EcSSB-GFP and DNA

We previously demonstrated that ssDNA exerts an inhibitory effect on EcSSB condensation *in vitro* ^9^. This observation raised the possibility that bacterial DNA (nucleoid regions) and EcSSB are spatially segregated in the absence of stress. To test this proposition and define the spatial relationship between DNA and EcSSB assemblies in cells, we used Hoechst 33342 DNA dye and analyzed cells during exponential and stationary growth phases (**Fig. 4; see Methods for details**). We observed limited overlap between EcSSB-GFP and Hoechst signals with a more pronounced separation in stationary-phase cells. These observations are consistent with spatial segregation between EcSSB-enriched regions and nucleoid DNA during periods of reduced DNA metabolic activity.

**Figure 4.**
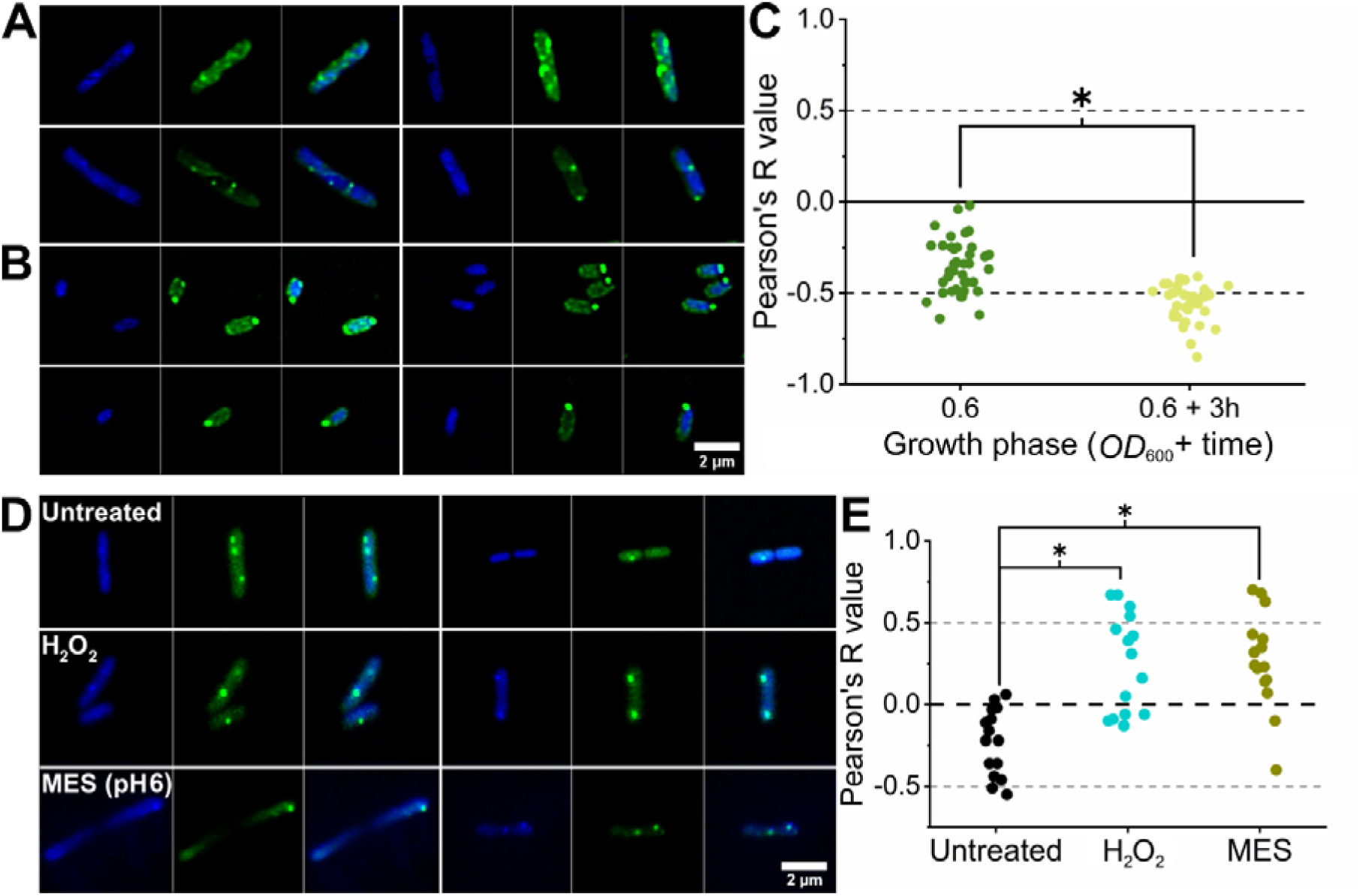
Spatial relationship between EcSSB-GFP and cellular DNA under physiological conditions and stress. Shown are SIM images of Hoechst 33342-stained (blue) E. coli #29003 cells (EcSSB-GFP, green) in the **(A)** exponential (OD_600_ = 0.6) and **(B)** stationary (OD_600_ = 0.6 + 3 h) growth phases. **(C)** Pearson’s analysis of EcSSB-GFP and Hoechst signals shows limited spatial overlaps in the exponential phase, which is further reduced in the stationary phase (Mann-Whitney test, p < 0.05). No preferential accumulation of DNA signal within EcSSB-GFP–positive regions was observed in either condition. **(D)** Confocal microscopy images showing Hoechst 33342-stained (blue) E. coli #29003 cells (EcSSB-GFP, green) in the exponential growth phase (OD_600_ = 0.6) in the absence and presence of stressors (see **Table S2** for conditions). **(E)** Pearson’s analysis of EcSSB-GFP and Hoechst signals shows significantly increased spatial overlap between EcSSB-GFP and DNA signals upon stress treatment (Mann-Whitney test, p < 0.05). In stressed cells, DNA signal was often observed in regions with stronger EcSSB-GFP fluorescence. This behavior was not detectable in untreated exponential-phase cells. Cells with H_2_O_2_ and MES treatment contained a smaller number of foci compared to stress-free cells, as was also seen in **Fig. 2**.

Upon treatment with H_2_O_2_ and MES (pH 6) (selected based on the above results as effective stressors), EcSSB-GFP foci were reduced, and a concomitant increase in spatial overlap between EcSSB-GFP and DNA signals was observed (**Fig. 4**). This effect was accompanied by the redistribution of EcSSB-GFP signal and reduced abundance of discrete foci. Overall, these results indicate stress-associated changes in the spatial relationship between EcSSB and DNA *in vivo*, consistent with our *in vitro* observations on EcSSB condensation behavior ^9^.

### *In vitro* assays support the physicochemical plausibility of EcSSB liquid condensate based cellular regulation

In addition to our *in vivo* experiments (**Figs. 1-4 and S1-3**), we carried out a comprehensive array of well-controlled *in vitro* biophysical measurements on EcSSB condensates to provide a quantitative framework connecting droplet-level morphologies in isolated molecular systems with cellular organization (**Figs. S4-S9**). These *in vitro* fluorescence microscopy, turbidity-based, microscale thermophoresis (MST), microfluidic resistive pulse sensing (MRPS), and dynamic light scattering (DLS) experiments support the view that (*i*) EcSSB condensation can occur when protein concentrations exceed a critical LLPS threshold which is lower than intracellular EcSSB concentration, (*ii*) cellular EcSSB-GFP foci may arise, at least in part, from the intrinsic condensation propensity of EcSSB, and (*iii*) the applied stressors modulate cellular EcSSB assemblies *in vivo* through cellular response mechanisms rather than through direct effects on EcSSB condensation.

First, we assessed whether EcSSB-GFP, expressed in the #29003 cell line applied in our experiments *in vivo*, can freely mix and form co-condensates with WT EcSSB **(Fig. S4)**. These experiments revealed that EcSSB-GFP containing condensates are present in #29003 cell extracts and they can be readily diluted with purified WT EcSSB, with no tendency for aggregation, demonstrating the utility of the EcSSB-GFP construct for *in vivo* visualization of EcSSB dynamics.

Second, we probed the chemical sensitivity of EcSSB droplets to condensation effectors **(Fig. S5)**. We found that the aliphatic alcohol 1,6-hexanediol, a well-established LLPS inhibitor ^34^, showed characteristic inhibition of EcSSB droplet formation at high concentrations (> 300 mM). The effective range of 1,6-hexanediol was similar to that observed earlier for condensation of the human SSB2 homolog ^35^. Phase-separating proteins have been shown to effectively form condensates under conditions with reduced water potential ^21^. Therefore, we assessed EcSSB condensation at increasing concentrations of sorbitol, an agent suitable to monitor LLPS under a broad range of osmotic conditions. EcSSB underwent phase separation even at 500 mg/ml sorbitol (**Fig. S5C-D**), which exceeds the macromolecule concentration in an osmotically stressed bacterial cytoplasm ^36^, indicating that EcSSB may retain the ability to undergo condensation under osmotically stressed conditions. The reduction in EcSSB condensation at extremely high sorbitol concentrations (> 500 mg/ml) probably originated from the increased viscosity of the medium (**Fig. S5C-D**). Moreover, we found that the organic solvent DMSO exhibited notable inhibitory effects on EcSSB condensation only at high concentrations (> 8 v/v%), highlighting the resilient nature of EcSSB condensates in the presence of organic solvents (**Fig. S5E-F**). This result also verifies that DMSO, as a small molecule solvent, is suitable for future applications screening small-molecule active substances for effects on EcSSB condensation.

Next, we performed *in vitro* microscale thermophoresis (MST) experiments to describe the concentration-dependent onset and time scale of EcSSB condensate formation **(Fig. S6)**. These measurements revealed that the critical EcSSB concentration for phase transition (2.5 µM) is below the physiological EcSSB concentration **(Fig. S6A-B)** ^37^. Moreover, we found that the condensation response of EcSSB is rapid (extensive condensation was already observed at 2 minutes after reaction onset), and the data reflected subsequent particle growth in the minutes time scale **(Fig. S6C-D)**. The NaCl concentration-dependent inhibition profile of the MST response was in line with the well-established chloride inhibition feature of EcSSB condensation ^9–11,30^, corroborating that the MST curves monitor *bona fide* protein LLPS processes **(Fig. S6E-F)**.

We defined the size distribution of individual EcSSB condensate particles *in vitro* by microfluidic resistive pulse sensing (MRPS) experiments **(Fig. S7)**. These measurements revealed lognormal size distributions for EcSSB condensates that scaled with protein concentration. This behavior provides additional evidence for the liquid nature of EcSSB condensates, while the condensate size scaling defined *in vitro* also parallels focus size variabilities observed *in vivo* (**Figs. 1-4**). In addition, our time-dependent dynamic light scattering (DLS) measurements of the hydrodynamic radius of EcSSB condensates provided evidence for rapid formation of EcSSB condensates with moderate growth on longer time scales **(Fig. S8A)**. We found that polydispersity decreases with EcSSB concentration, reinforcing condensate maturation/coalescence occurring at higher EcSSB concentrations **(Fig. S8B)**, in line with EcSSB-GFP foci dynamics observed *in vivo* across growth phases (**Fig. 1**). Together, the above *in vitro* findings (**Figs. S4-S8**) support a physicochemical framework for EcSSB condensation that mirrors the behavior of previously observed *in vivo* foci across growth phases and under stress (**Figs. 1-4**).

### Testing the effects of *in vivo* stressors on EcSSB condensates *in vitro*

To define the direct effects of the stressors applied *in vivo* (**Figs. 2-4**) on EcSSB condensation *in vitro*, turbidity assays (λ = 600 nm) were conducted alongside epifluorescence imaging of purified EcSSB samples **(Fig. S9)**. The results revealed that most of the applied stressors do not influence the formation of EcSSB condensates *in vitro*, even at high stressor concentrations. Notably, despite the previously defined inhibitory effect of NaCl on EcSSB condensation *in vitro* ^9,11^, this agent even increased foci formation *in vivo* (**Fig. 2**). Besides NaCl, L-Arg was the only effector that exhibited marked EcSSB condensation inhibition *in vitro* at the dose applied *in vivo*. The presence of spherical, phase-separated droplets in epifluorescence microscopy images demonstrates that the turbidity signals originate from LLPS rather than amorphous protein aggregation in the case of most applied stressors. Assuming that the observed EcSSB-GFP foci represent EcSSB condensates *in vivo*, these data suggest that the applied stressors modulate them indirectly through cellular response mechanisms rather than via direct physicochemical effects.

## Discussion

### The cellular EcSSB pool is distributed between genome-associated complexes and storage assemblies with condensate properties

In this work, we combined super-resolution microscopy with quantitative image analysis to determine the spatial organization of EcSSB *in vivo*. We found, using an EcSSB-GFP construct, that EcSSB typically forms multiple intracellular assemblies under stress-free and exponential growth conditions, which are spatially separated from DNA to a large extent (**Figs. 1** and **4**). Notably, stationary-phase cells, characterized by reduced proliferation, contain EcSSB assemblies that are brighter but smaller in number, with even more pronounced separation from DNA (**Figs. 1** and **4**). Various stressors and metabolic activation markedly reduce the abundance of EcSSB-containing assemblies in exponentially growing cells, while corresponding responses were attenuated in stationary phase. Characteristically, these events also trigger increased association of EcSSB-GFP foci with DNA signal (**Figs. 2** and **4**). These observations, together with EcSSB’s LLPS propensity that is abolished by ssDNA binding ^9,10^, are consistent with the cellular EcSSB pool being distributed between replication- and repair-associated, DNA-bound EcSSB complexes ^6,23,25,26,31^ and cytoplasmic storage assemblies. The material properties of the latter remain to be precisely defined. However, besides the physical properties of EcSSB LLPS droplets *in vitro* (**Figs. S4-S8**) ^9,10^, several lines of evidence point to the existence and liquid condensate nature of cellular EcSSB storage pools, as discussed below.

First, the size and fluorescence intensity of cellular EcSSB assemblies are substantially greater than that expected for individual replication forks. Based on previous estimates, replication forks in *E. coli* typically expose approximately 0.5–1 kb of ssDNA during active replication. Assuming an average footprint of ∼35–40 nucleotides per EcSSB tetramer, this corresponds to roughly 15–25 tetramers per replication fork, in line with 5-11 EcSSB tetramers per replisome calculated from *in vivo* single-molecule fluorescence microscopy data ^24,38^. In the current study we observed EcSSB foci with diameters typically ranging from 100–500 nm. Based on the crystal structure of EcSSB (PDB: 4MZ9) in which EcSSB forms a tetrameric assembly with a ∼50 Å diameter, a few thousand tetramers packed in a dense molecular packing would result in such foci sizes. We note, however, that this estimate does not account for differences in packing density between crystals and LLPS droplets, the structural effects of EcSSB’s intrinsically disordered tail region, and the possible incorporation of EcSSB’s interacting partners, all of which could influence condensate size *in vivo.* Moreover, our fluorescence intensity analysis revealed that up to 40% of the total EcSSB-GFP signal was concentrated in foci during exponential growth, whereas this proportion increased to 70% in stationary-phase cells. Bonde *et al.* found that *E. coli* cells expressing only a variant of EcSSB lacking the IDL (but retaining the CTP, SSBΔ^120–166^), which is known to be less prone to undergo LLPS ^10^, contained about half the amount of EcSSB compared to the WT strain (∼8800 ± 2700 monomers of SSBΔ^120–166^ *versus* 19700 ± 5600 monomers of WT per cell) ^5^. Although these measurements do not directly establish the nature or composition of the observed foci, they quantitatively support the existence of EcSSB storage assemblies of significant size.

Second, in accordance with the results of Zhao *et al*., we frequently observed EcSSB assemblies in peripheral regions of the cytoplasm with little spatial overlap with DNA signal during stationary phase in which replication activity is substantially reduced (**Fig. 4**) ^29^. Moreover, a previous study revealed EcSSB assemblies even prior to replication initiation ^24^, while others observed only partial colocalization of EcSSB assemblies with replication fork-associated DnaQ ^26^ or with the nucleoid ^6^. These findings further point to the existence of non-DNA-associated EcSSB pools.

Third, we observed stress-induced EcSSB remodeling and increased DNA association under stress conditions (*e.g.* H_2_O_2_ and MES pH 6) that affect cellular physiology beyond DNA replication (**Fig. 4**). These findings are also consistent with stress-induced EcSSB mobilization upon the appearance of increased amounts of ssDNA, consistent with EcSSB condensate behavior *in vitro* whereby ssDNA binding competes with multivalent interactions driving EcSSB phase separation ^9,11,30^. The dynamic nature of intracellular SSB rearrangements was also demonstrated by FRAP experiments showing exchange times in the seconds range ^24,25^. In addition, the finding that SSB exchange is slowed down by hydroxyurea treatment is also consistent with SSB recruitment from cytoplasmic condensate pools to sites of active replication ^24^. Notably, NaCl stress enhanced EcSSB-GFP foci formation *in vivo*, despite chloride ions being strong *in vitro* inhibitors of EcSSB condensation ^9,11,30^. NaCl presumably causes osmotic changes in the cytoplasm, which could increase the local concentration of EcSSB, thereby driving condensate formation inside the cell (**Figs. 2-4**). In addition, we observed the disappearance of EcSSB-GFP foci following UV exposure (**Fig. 2**). This finding is in line with observations of Cherry *et al.* who classified EcSSB-mTur2 structures into bright, discrete puncta; broader EcSSB clusters; and diffuse signals. Following UV treatment, they likewise observed a reduction in the number of puncta and clusters, followed by their gradual reappearance over time, with clusters becoming the predominant species during recovery ^26^. The differential response of these assemblies to UV-induced DNA damage likely reflects differences in their underlying molecular organization and biological roles. Another observation with relevance to condensation-driven organization is the disappearance of EcSSB foci at elevated temperature ^23^. Although multiple physiological factors are altered under heat stress, this behavior is consistent with the pronounced temperature dependence of EcSSB condensation *in vitro* ^11,30^.

Taken together, the above discussed observations strongly suggest that EcSSB is distributed between DNA-associated complexes and cytoplasmic storage (liquid condensate or membrane bound ^29^) pools, which interchange *via* dynamic mobilization upon stress or physiological changes.

### Functional implications of intracellular EcSSB condensation

Recent discoveries on liquid-liquid phase separation (LLPS) have transformed our understanding of cellular organization in eukaryotes. In contrast, the role of LLPS in bacteria remains poorly understood, despite growing evidence suggesting that phase transition processes play central roles in bacterial stress adaptation and post-stress resuscitation ^17,39^. How bacterial cells detect and respond to stress—and whether LLPS constitutes a mechanistic basis for these processes—remains a fundamental scientific question.

Results from previous *in vitro* studies suggested a potential mechanism for EcSSB condensate dissolution. ssDNA was shown to become enriched within EcSSB condensates and bind to the tetramers. This binding, however, competes with the multivalent protein-protein interactions that drive EcSSB phase separation. Once the ssDNA concentration reaches a critical threshold, this competition is sufficient to initiate condensate dissolution, ultimately leading to complete disassembly ^9,11,30^. In the cytoplasm, this process may be promoted by increasing concentrations of genomic ssDNA stretches during stress or elevated metabolic activity. This interpretation is further supported by our *in vivo* observations showing limited colocalization between DNA and EcSSB-GFP foci under basal conditions, whereas stress conditions resulted in increased association of DNA with EcSSB-GFP foci (**Fig. 4**).

In addition to the above considerations, we previously showed that RecQ helicase, one of EcSSB’s key interacting partners, becomes enriched within EcSSB condensates *in vitro*, suggesting co-sequestration of genome maintenance factors *in vivo* ^9,26^. Thus, the loss of EcSSB’s LLPS propensity could also affect the availability of other DNA maintenance proteins. Moreover, these associated proteins are presumably released in a coordinated manner with EcSSB upon stress-induced dissolution of condensates. Disruption of EcSSB-driven spatial organization could impair this coordinated release and, consequently, compromise the efficiency of DNA repair processes. In addition, sequestration could protect sensitive factors from degradation, reducing energy consumption and enhancing survival.

Numerous adaptive bacterial systems support long-term cell growth and survival primarily under fluctuating environmental conditions and stress exposure; therefore, such selective advantages may not be readily apparent in standard growth assays ^39–42^. Bonde *et al.* found no severe growth defects in *E. coli* cells expressing previously described EcSSB variants with potentially impaired condensation capacity ^10^ under nutrient-rich laboratory conditions ^5^. Notably, however, the same study reported pronounced UV sensitivity of cells expressing the SSBΔ^120–166^ and SSBΔ^130–166^ mutants, which lack most of their IDLs (but not the CTP) and therefore are expected to be defective for phase separation but can bind partners. Moreover, in the same study, cells expressing SSBΔ^120–166^ exhibited a strong competitive disadvantage, despite showing growth kinetics comparable to wild-type cells in monoculture conditions ^5^. These results are consistent with EcSSB LLPS conferring a selective advantage under real-life competitive and stress-involving conditions ^30^. In addition, compromised EcSSB condensation propensity could affect genome stability, as supported by previous findings showing that cells expressing EcSSB tetramers with only two IDLs, presumably with reduced LLPS propensity and partner binding capacity, exhibit increased mutation rates ^43^. Nonetheless, an elevated mutational burden could, under certain conditions such as antibiotic exposure, facilitate the emergence of resistance to antimicrobial agents over shorter timescales. Testing these hypotheses and determining the phenotypic consequences of altered condensation propensity will require extended investigations using genetically engineered EcSSB variants that are incapable of undergoing LLPS while retaining their ssDNA-binding function ^30^.

Beyond stress adaptation, the observations presented here may support an emerging view of prokaryotic material organization mechanisms involving dynamic protein condensation ^21,44–46^. Global cytoplasmic transitions in bacteria from fluid-like to glass-like states have been reported in response to decreased metabolic activity ^21^, and metabolism-driven cytoplasmic fluidization has been linked to enhanced molecular mobility ^21,47^. Our findings on growth phase and metabolism-dependent EcSSB organization suggest that this mechanism may contribute to these broader cytoplasmic phase transitions during metabolic shifts and adaptation. Together with previous findings, the current work points to EcSSB assemblies being a dynamic and tunable component of bacterial DNA metabolic regulation (**Fig. 5**). Hence, these insights may also open avenues for new-mechanism antimicrobial discovery by targeting phase separation pathways.

**Figure 5.**
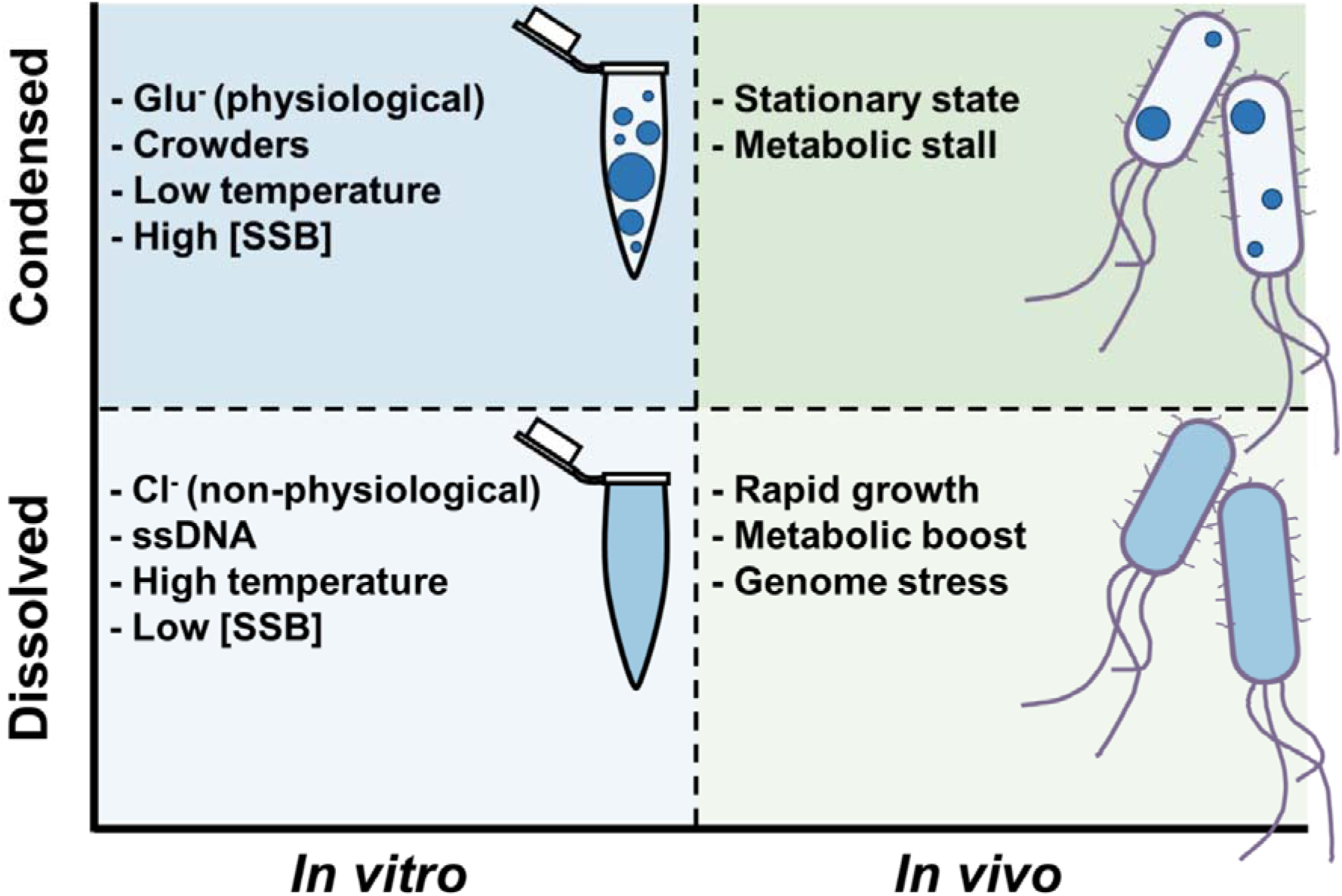
“Phase map” of EcSSB. EcSSB condensates are highly dynamic structures whose formation and dissolution are regulated by multiple factors in vitro ^9,11,30^ and possibly in vivo (this paper). This responsiveness enables EcSSB condensates to function as efficient tools for bacterial cells, facilitating the sequestration of proteins when not required and enabling their rapid mobilization to sites of action upon demand. Note that in vitro conditions mimicking the unstressed cellular environment promote condensation.

## Materials and Methods

### Materials

Stressors used in this study, including their applied dose and mechanism of action, are described in detail in **Table S2**. Recombinant EcSSB expression and purification, as well as AF555 (AlexaFluor 555) and IAF (5-iodoacetamidofluorescein) labeling was carried out as described in Harami *et al*. ^9^. Since EcSSB does not contain any cysteines, the EcSSB^G26C^ variant was used for IAF labeling. AF555 labeling was performed on wild type EcSSB. EcSSB protein concentrations are expressed in terms of EcSSB tetramers throughout this article.

### Cell culture preparation for fluorescence imaging

Cells were grown overnight in Luria–Bertani medium (LB) containing kanamycin (25 µg/ml) and chloramphenicol (34 µg/ml) at 37°C. Overnight cultures were diluted 1:100 in fresh LB media containing antibiotics and grown at room temperature at 200 rpm shaking until they reached a certain *OD*_600_ value + time indicating the different growth phases (exponential, stationary or death phase). We note that cells were grown under these conditions to improve EGFP signal detection during imaging. Cells were centrifuged, washed with 10 mM MgSO_4_ twice, then resuspended in MgSO_4_ containing the stressors at concentrations indicated in **Table S2**. The pH of stressor solutions was adjusted to 7.0 except for MES (pH 6) when necessary. Cells were shaken for 30 minutes at 200 rpm at 37 °C and subsequently imaged without any fixation or L-lysine treatment. Depending on cell density, dilutions were applied before imaging to avoid a confluence of cells during imaging. Notably, when cells in different growth phases were studied, no incubation in MgSOO was used, and they appeared identical to the untreated cells in the stress panels in which cells were in MgSO_4_ for 30 minutes. This indicates that a 30-minute incubation in MgSOO does not significantly alter EcSSB organization patterns. UV-stressed cells were also observed without incubation to avoid regeneration.

When cell lines #29003 and #27869 were compared in **Fig. S1**, cells were incubated with 2 µM FM 4-64 membrane dye in 10 mM MgSO_4_ for 2 minutes. Cells were washed twice before imaging.

### Super-resolution (SIM) and confocal microscopy of *E. coli* cells

For SIM microscopy, Zeiss High Performance 0.17 ± 0.005 mm thick glass coverslips and a Zeiss Elyra S1 SIM microscope with 488-nm (Lasos) laser were used. Hoechst 33342 (Biotium, #40046) DNA stain images were captured using a Nikon X-Light V3 Deep SIM microscope (LED, solid state from Lumencor). Cells were treated with Hoechst for one minute followed by two wash steps before observation. No noticeable changes in cell morphology and EcSSB patterns were observed between stained and unstained cells (*cf.* **Figs. 1** and **4**), demonstrating that the cells remained viable upon staining.

For confocal microscopy, a Zeiss LSM710 microscope was used with Menzel-Glaser (20x20 mm, Thermo scientific) glass cover slides. Hoechst 33342 staining was performed as described for SIM imaging.

### Evaluation of microscopy images of *E. coli* cells

Image analysis was performed using Fiji software. To ensure accurate quantification, the brightest foci were checked for signal saturation, and no evidence of saturation or clipping artifacts was found. For image evaluation purposes, two scripts were developed for a more automatic quantification of protein assemblies in cells **(Supplementary file 1)**. During evaluation, cell segmentation was performed using Li cross-entropy thresholding, which determines the optimal intensity cutoff by minimizing cross-entropy. This method is robust for asymmetric and noisy intensity distributions, making it well-suited for automatic detection of cells in fluorescence images. This step was followed by another thresholding step within the cells to identify foci using a modified Find Maxima function (**Fig. S10, Supplementary file 1**). A 2D Gaussian was fitted to each spot, and intensities were calculated by measuring the space under the Gaussian curve and correcting with the background at the base of the curves. The script can be fine-tuned at this step, as the Find Maxima function allows adjustment of the minimum intensity required for spot detection relative to the background. For the current study, the parameters were set based on untreated cells and applied identically to all treated samples to allow direct comparison. To determine the percentage of fluorescent signal present in the foci compared to the cytoplasm, the developed script (**Supplementary file 1**) was used to outline foci and to subtract them from the original image. The difference between intensities was used to define material percentage within foci (**Fig. S2**). Although bacteria may exhibit autofluorescence in the GFP channel, this contribution was expected to be comparable across samples. To minimize the effect of background fluorescence, images were background-subtracted prior to fluorescence quantification and ratio calculations. Colocalization analysis of images of Hoechst-stained cells was performed using Fiji plugin Coloc2. For colocalization analysis, EcSSB-GFP foci were first identified and segmented using the GFP channel. Regions of interest (ROIs) corresponding to individual EcSSB assemblies were then used to quantify the spatial relationship between EcSSB and DNA signals. Colocalization metrics were calculated within these segmented focal regions rather than across the entire cellular area, thereby reducing biases arising from the sparse and heterogeneous distribution of nucleoid-associated DNA and GFP signals. To test the effect of the FM 4-64 membrane dye at different concentrations, the dye was applied at 2 µM and 8 µM (**Fig. S11**). In addition, we compared unprocessed and processed SIM images to quantify the gain in resolution provided by the SIM technique (**Fig. S12**).

### Growth assays

To investigate the effects of EcSSB-GFP expression on *E. coli*, the growth of strains #29003 and #27869 was monitored in LB medium for 24 h, both in the absence and presence of the indicated stressors. Stressors were applied at the concentrations listed in **Table S2**. Growth assays were performed in 96-well microplates (VWR Tissue Culture Plate). Each well was inoculated with 6 × 10O cells in the exponential growth phase (*OD*_600_ = 0.6), and cell growth was monitored by measuring *OD*OO O every 10 min using a Tecan Infinite F Nano+ plate reader following 10 s of orbital shaking at 200 rpm.

### General measurement conditions for *in vitro* experiments

*In vitro* measurements were performed at 25°C in LLPS buffer containing 20 mM Tris-acetate (Tris-OAc) pH 7.5, 5 mM MgOAc and 50 mM NaGlu.

### Turbidity assays of purified EcSSB condensates

Turbidity measurements were performed at 600 nm, using Thermo Fisher Nunc 384-well, non-treated flat bottom microplates and a Tecan Infinite F Nano+ plate reader instrument. The sample volume was 100 µL, in a buffer containing 20 mM Tris-OAc pH 7.5, 5 mM MgOAc and 50 mM NaGlu. The EcSSB tetramer concentration was 20 µM. Measurements were performed in triplicate at room temperature.

### Enrichment of EcSSB-GFP in EcSSB LLPS condensates

*E. coli* #29003 cells were grown in LB medium with shaking at 200 rpm. Cells were harvested by centrifugation (7,459 × *g*, 10 min, 4°C) and resuspended in 35 mL of 50 mM Tris-HCl (pH 8.0), 150 mM NaCl buffer. Cells were lysed by ultrasonication, and the lysate was clarified by centrifugation (48,384 × *g*, 20 min, 4°C). The resulting supernatant was diluted at a 1:5 ratio in LLPS buffer containing 3% PEG 20,000, and the GFP signal was monitored by fluorescence microscopy (Nikon Eclipse Ti-E TIRF, similarly as described in the next section). The same dilution and imaging procedure was performed in the presence of either 5 µM purified EcSSB tetramer ^30^ or EGFP.

### Epifluorescence microscopy of purified EcSSB condensates

The effects of stressors on EcSSB condensates *in vitro* were visualized using epifluorescence microscopy. Samples contained 20 µM EcSSB tetramers and 0.3 µM fluorescein-labeled EcSSB, which readily became enriched in EcSSB droplets ^9^. Samples were measured on Ibidi

15 Well µ-Slide Angiogenesis slides at room temperature. A Nikon Eclipse Ti-E TIRF microscope was used in epifluorescence mode with a Cyan 488-nm laser (Coherent). Determination of total droplet area in middle ROI (area: 600 x 600 pixels; X, Y coordinates from left, uppermost pixel position: 300, 300 pixels) was carried out as in ^48^.

### Microscale thermophoresis (MST)

The thermophoretic mobility of 5 µM EcSSB (containing 50 nM iodoacetamidofluorescein (IAF)-labeled EcSSB^G26C^) was monitored using a Nanotemper Monolith device (set to 20 % MST power and 75 % excitation power) in LLPS buffer. Premium capillaries were used for sample loading. Capillary scans showed ideal capillary shapes and no protein adhesion to the surface. Baseline was recorded for 5 seconds before infrared irradiation for 30 seconds. Fluorescence recovery was monitored for additional 5 seconds after the end of irradiation. Fluorescence was normalized to baseline. Thermophoretic mobility was determined as the fraction of the baseline fluorescence signal (*F*_norm_) measured at 20 seconds irradiation.

### Microfluidic resistive pulse sensing (MRPS)

MRPS measurements were performed using an nCS1 instrument (Spectradyne LLC, USA). Data obtained were analyzed using Viewer software (Spectradyne LLC, USA). EcSSB was diluted in LLPS buffer unless otherwise indicated. Samples were measured using a C-2000 cartridge (∼250Onm to 2000Onm measurement range).

### Dynamic light scattering (DLS)

EcSSB samples were diluted in LLPS buffer and measured in low volume disposable cuvettes with 1 cm path-length (UVette, Eppendorf Austria GmbH) using a W130i dynamic light scattering device (DLS, Avid Nano Ltd., High Wycombe, UK) equipped with a diode laser (660 nm) and a photodiode detector.

## Data Availability

The data underlying this article are available in the article and in its online Supplementary Material. The data underlying this article will be shared upon reasonable request to the corresponding author.

## Supplementary Data

Supplementary Data are provided with this article.

## Author Contributions

**P.E.**: Conceptualization, Data curation, Formal Analysis, Investigation, Methodology Validation, Visualization, Writing – original draft, Writing – review & editing, Funding acquisition. **J.S.**: Investigation, Data curation, Validation, Writing – original draft. **J.P.**: Investigation, Methodology. **B.J.**: Conceptualization, Methodology. **V.K.**: Investigation. **Z.J.K.**: Investigation. **H.H.**: Investigation, Methodology. **T.S.**: Investigation. **T.J.**: Investigation, Methodology. **T.B.S.**: Conceptualization, Methodology. **S.B.**: Investigation, Methodology. **E.S.M.**: Investigation, Methodology. **M.K.**: Conceptualization, Funding acquisition, Project administration, Resources, Supervision, Writing – review & editing. We thank Gábor M. Harami for insightful discussions in the topic of the manuscript.

## Supporting information

Scripts

## References

1 Chatterjee, N. & Walker, G. C. Mechanisms of DNA damage, repair, and mutagenesis. Environmental and molecular mutagenesis 58, 235–263, doi:10.1002/em.22087 (2017).

2 Shereda, R. D., Kozlov, A. G., Lohman, T. M., Cox, M. M. & Keck, J. L. SSB as an organizer/mobilizer of genome maintenance complexes. Critical reviews in biochemistry and molecular biology 43, 289–318, doi:10.1080/10409230802341296 (2008).

3 Bonde, N. J., Kozlov, A. G., Cox, M. M., Lohman, T. M. & Keck, J. L. Molecular insights into the prototypical single-stranded DNA-binding protein from E. coli. Critical reviews in biochemistry and molecular biology 59, 99–127, doi:10.1080/10409238.2024.2330372 (2024).

4 Raghunathan, S., Kozlov, A. G., Lohman, T. M. & Waksman, G. Structure of the DNA binding domain of E. coli SSB bound to ssDNA. Nature structural biology 7, 648–652, doi:10.1038/77943 (2000).

5 Bonde, N. J., Henry, C., Wood, E. A., Cox, M. M. & Keck, J. L. Interaction with the carboxy-terminal tip of SSB is critical for RecG function in E. coli. Nucleic acids research 51, 3735–3753, doi:10.1093/nar/gkad162 (2023).

6 Dubiel, K. et al. Development of a single-stranded DNA-binding protein fluorescent fusion toolbox. Nucleic acids research 48, 6053–6067, doi:10.1093/nar/gkaa320 (2020).

7 Antony, E. & Lohman, T. M. Dynamics of E. coli single stranded DNA binding (SSB) protein-DNA complexes. Seminars in cell & developmental biology 86, 102–111, doi:10.1016/j.semcdb.2018.03.017 (2019).

8 Shereda, R. D., Reiter, N. J., Butcher, S. E. & Keck, J. L. Identification of the SSB binding site on E. coli RecQ reveals a conserved surface for binding SSB’s C terminus. Journal of molecular biology 386, 612–625, doi:10.1016/j.jmb.2008.12.065 (2009).

9 Harami, G. M. et al. Phase separation by ssDNA binding protein controlled via protein-protein and protein-DNA interactions. Proceedings of the National Academy of Sciences of the United States of America 117, 26206–26217, doi:10.1073/pnas.2000761117 (2020).

10 Kozlov, A. G. et al. How Glutamate Promotes Liquid-liquid Phase Separation and DNA Binding Cooperativity of E. coli SSB Protein. Journal of molecular biology 434, 167562, doi:10.1016/j.jmb.2022.167562 (2022).

11 Kovacs, Z. J. et al. Fine-tuned interactions between globular and disordered regions of single-stranded DNA binding (SSB) protein are required for dynamic condensation under physiological conditions. Protein science : a publication of the Protein Society 34, e70109, doi:10.1002/pro.70109 (2025).

12 Guo, D. et al. Liquid-Liquid phase separation in bacteria. Microbiological research 281, 127627, doi:10.1016/j.micres.2024.127627 (2024).

13 Uversky, V. N. Biological Liquid-Liquid Phase Separation, Biomolecular Condensates, and Membraneless Organelles: Now You See Me, Now You Don’t. International journal of molecular sciences 24, doi:10.3390/ijms241713150 (2023).

14 Tong, X. et al. Liquid-liquid phase separation in tumor biology. Signal transduction and targeted therapy 7, 221, doi:10.1038/s41392-022-01076-x (2022).

15 Peng, P. H., Hsu, K. W. & Wu, K. J. Liquid-liquid phase separation (LLPS) in cellular physiology and tumor biology. American journal of cancer research 11, 3766–3776 (2021).

16 Wang, B. et al. Liquid-liquid phase separation in human health and diseases. Signal transduction and targeted therapy 6, 290, doi:10.1038/s41392-021-00678-1 (2021).

17 Jin, X. et al. Membraneless organelles formed by liquid-liquid phase separation increase bacterial fitness. Science advances 7, eabh2929, doi:10.1126/sciadv.abh2929 (2021).

18 Gomes, E. & Shorter, J. The molecular language of membraneless organelles. The Journal of biological chemistry 294, 7115–7127, doi:10.1074/jbc.TM118.001192 (2019).

19 Poudyal, M. et al. Intermolecular interactions underlie protein/peptide phase separation irrespective of sequence and structure at crowded milieu. Nature communications 14, 6199, doi:10.1038/s41467-023-41864-9 (2023).

20 Feng, Z., Chen, X., Wu, X. & Zhang, M. Formation of biological condensates via phase separation: Characteristics, analytical methods, and physiological implications. The Journal of biological chemistry 294, 14823–14835, doi:10.1074/jbc.REV119.007895 (2019).

21 Azaldegui, C. A., Vecchiarelli, A. G. & Biteen, J. S. The emergence of phase separation as an organizing principle in bacteria. Biophysical journal 120, 1123–1138, doi:10.1016/j.bpj.2020.09.023 (2021).

22 Alberti, S., Gladfelter, A. & Mittag, T. Considerations and Challenges in Studying Liquid-Liquid Phase Separation and Biomolecular Condensates. Cell 176, 419–434, doi:10.1016/j.cell.2018.12.035 (2019).

23 Reyes-Lamothe, R., Possoz, C., Danilova, O. & Sherratt, D. J. Independent positioning and action of Escherichia coli replisomes in live cells. Cell 133, 90–102, doi:10.1016/j.cell.2008.01.044 (2008).

24 Reyes-Lamothe, R., Sherratt, D. J. & Leake, M. C. Stoichiometry and architecture of active DNA replication machinery in Escherichia coli. Science 328, 498–501, doi:10.1126/science.1185757 (2010).

25 Spenkelink, L. M. et al. Recycling of single-stranded DNA-binding protein by the bacterial replisome. Nucleic acids research 47, 4111–4123, doi:10.1093/nar/gkz090 (2019).

26. Cherry, M. E. et al. Spatiotemporal Dynamics of Single-stranded DNA Intermediates in Escherichia coli. *bioRxiv*, 2023.2005.2008.539320, doi:10.1101/2023.05.08.539320 (2023).

27 Ripandelli, R. A. A., Wood, E. A., Robinson, A., van Oijen, A. M. & Cox, M. M. Spatiotemporal characterization of single-stranded DNA intermediates after UV irradiation: II. Rapid growth and effects of recA and recJ. PLoS genetics 22, e1012110, doi:10.1371/journal.pgen.1012110 (2026).

28 Sharma, N. et al. Spatiotemporal characterization of single-stranded DNA Intermediates after UV Irradiation: I: Post-replication gaps formed during slow growth. PLoS genetics 22, e1012109, doi:10.1371/journal.pgen.1012109 (2026).

29 Zhao, T. et al. Super-resolution imaging reveals changes in Escherichia coli SSB localization in response to DNA damage. Genes to cells : devoted to molecular & cellular mechanisms 24, 814–826, doi:10.1111/gtc.12729 (2019).

30 Ecsedi, P. et al. Selective engineering of condensation properties of single-stranded DNA binding (SSB) protein via its intrinsically disordered linker region. Nucleic acids research 53, doi:10.1093/nar/gkaf481 (2025).

31 Liu, J. et al. Novel, fluorescent, SSB protein chimeras with broad utility. Protein science : a publication of the Protein Society 20, 1005–1020, doi:10.1002/pro.633 (2011).

32 Kram, K. E., Henderson, A. L. & Finkel, S. E. Escherichia coli Has a Unique Transcriptional Program in Long-Term Stationary Phase Allowing Identification of Genes Important for Survival. mSystems 5, doi:10.1128/mSystems.00364-20 (2020).

33 Schink, S. J., Biselli, E., Ammar, C. & Gerland, U. Death Rate of E. coli during Starvation Is Set by Maintenance Cost and Biomass Recycling. Cell systems 9, 64–73 e63, doi:10.1016/j.cels.2019.06.003 (2019).

34 Duster, R., Kaltheuner, I. H., Schmitz, M. & Geyer, M. 1,6-Hexanediol, commonly used to dissolve liquid-liquid phase separated condensates, directly impairs kinase and phosphatase activities. The Journal of biological chemistry 296, 100260, doi:10.1016/j.jbc.2021.100260 (2021).

35 Kovacs, Z. J. et al. DNA-dependent phase separation by human SSB2 (NABP1/OBFC2A) protein points to adaptations to eukaryotic genome repair processes. Protein science : a publication of the Protein Society 33, e4959, doi:10.1002/pro.4959 (2024).

36 van den Bogaart, G., Hermans, N., Krasnikov, V. & Poolman, B. Protein mobility and diffusive barriers in Escherichia coli: consequences of osmotic stress. Molecular microbiology 64, 858–871, doi:10.1111/j.1365-2958.2007.05705.x (2007).

37 Bobst, E. V., Bobst, A. M., Perrino, F. W., Meyer, R. R. & Rein, D. C. Variability in the nucleic acid binding site size and the amount of single-stranded DNA-binding protein in Escherichia coli. FEBS letters 181, 133–137, doi:10.1016/0014-5793(85)81128-5 (1985).

38 Bianco, P. R. & Lu, Y. Single-molecule insight into stalled replication fork rescue in Escherichia coli. Nucleic acids research 49, 4220–4238, doi:10.1093/nar/gkab142 (2021).

39 Leplae, R. et al. Diversity of bacterial type II toxin-antitoxin systems: a comprehensive search and functional analysis of novel families. Nucleic acids research 39, 5513–5525, doi:10.1093/nar/gkr131 (2011).

40 Pizzolato-Cezar, L. R., Spira, B. & Machini, M. T. Bacterial toxin-antitoxin systems: Novel insights on toxin activation across populations and experimental shortcomings. Current research in microbial sciences 5, 100204, doi:10.1016/j.crmicr.2023.100204 (2023).

41 Nair, S. & Finkel, S. E. Dps protects cells against multiple stresses during stationary phase. Journal of bacteriology 186, 4192–4198, doi:10.1128/JB.186.13.4192-4198.2004 (2004).

42 Schellhorn, H. E. Function, Evolution, and Composition of the RpoS Regulon in Escherichia coli. Frontiers in microbiology 11, 560099, doi:10.3389/fmicb.2020.560099 (2020).

43 Antony, E. et al. Multiple C-terminal tails within a single E. coli SSB homotetramer coordinate DNA replication and repair. Journal of molecular biology 425, 4802–4819, doi:10.1016/j.jmb.2013.08.021 (2013).

44 Janissen, R. et al. Global DNA Compaction in Stationary-Phase Bacteria Does Not Affect Transcription. Cell 174, 1188–1199 e1114, doi:10.1016/j.cell.2018.06.049 (2018).

45 Calhoun, L. N. & Kwon, Y. M. Structure, function and regulation of the DNA-binding protein Dps and its role in acid and oxidative stress resistance in Escherichia coli: a review. Journal of applied microbiology 110, 375–386, doi:10.1111/j.1365-2672.2010.04890.x (2011).

46 Almiron, M., Link, A. J., Furlong, D. & Kolter, R. A novel DNA-binding protein with regulatory and protective roles in starved Escherichia coli. Genes & development 6, 2646–2654, doi:10.1101/gad.6.12b.2646 (1992).

47 Parry, B. R. et al. The bacterial cytoplasm has glass-like properties and is fluidized by metabolic activity. Cell 156, 183–194, doi:10.1016/j.cell.2013.11.028 (2014).

48 Harami, G. M. et al. Redox-dependent condensation and cytoplasmic granulation by human ssDNA-binding protein-1 delineate roles in oxidative stress response. iScience 27, 110788, doi:10.1016/j.isci.2024.110788 (2024).

