## Supplementary material for "Phase transition of bacterial single-stranded DNA binding (SSB) protein upon stress response and metabolic adaptation": SI

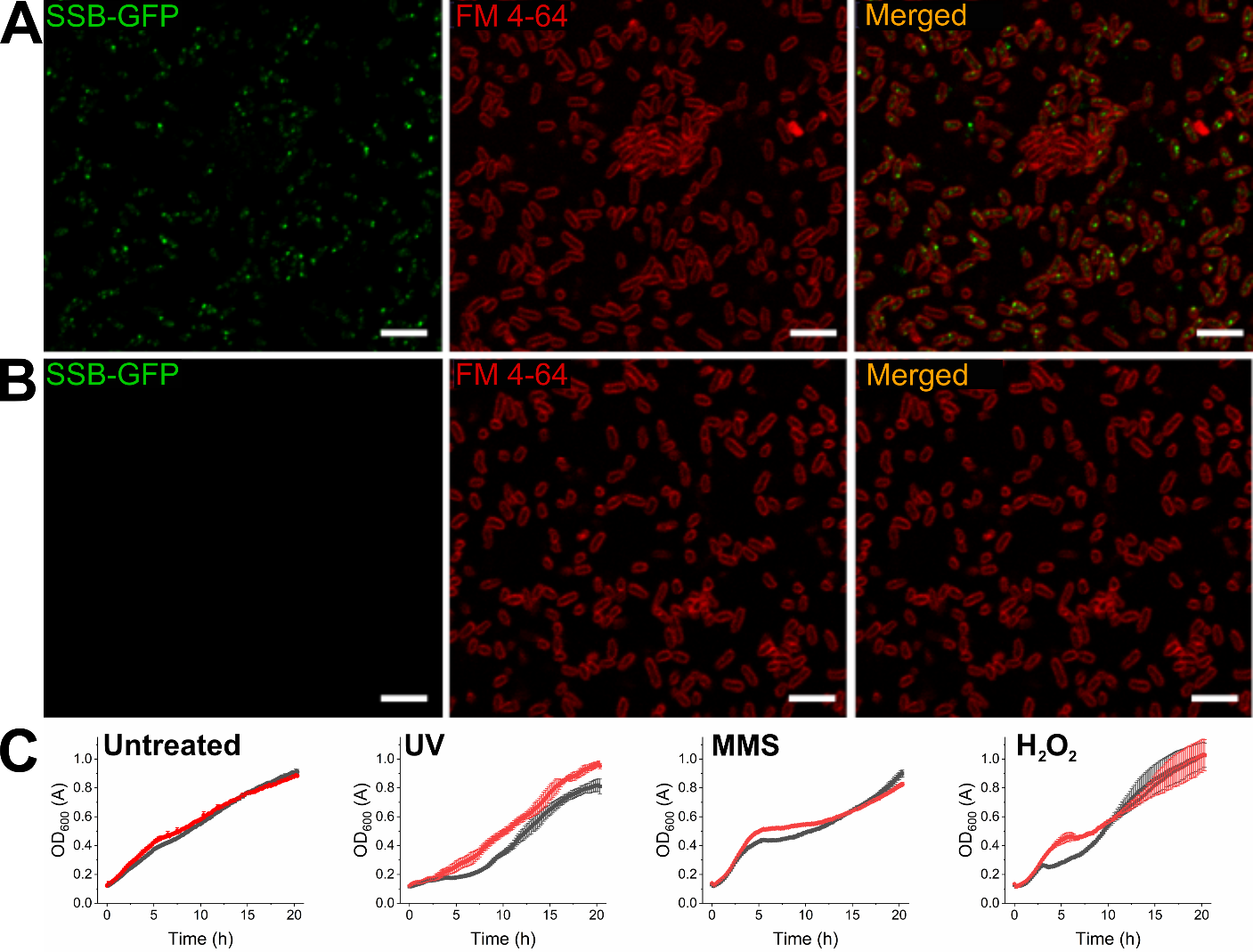

**Figure S1. Comparison of EcSSB-GFP expressing and non-expressing cells.** Shown are confocal images of cells in the exponential growth phase (**A**) expressing EcSSB-GFP (#29003) and (**B**) not expressing EcSSB-GFP (#27869) ^1^. Cells were stained with the membrane dye FM 4-64 (2 µM). The images show that EcSSB-GFP expression does not detectably affect cell morphology. (**C**) Cell growth at 25°C was monitored in cells expressing EcSSB-GFP (red) and non-expressing cells (black), both in the absence (left panel) and in the presence of the indicated stressors at the concentrations used for the *in vivo* measurements described in this study **(Table S2)**. Data represent mean ± SD of three independent replicates. Scale bars, 5 µm.

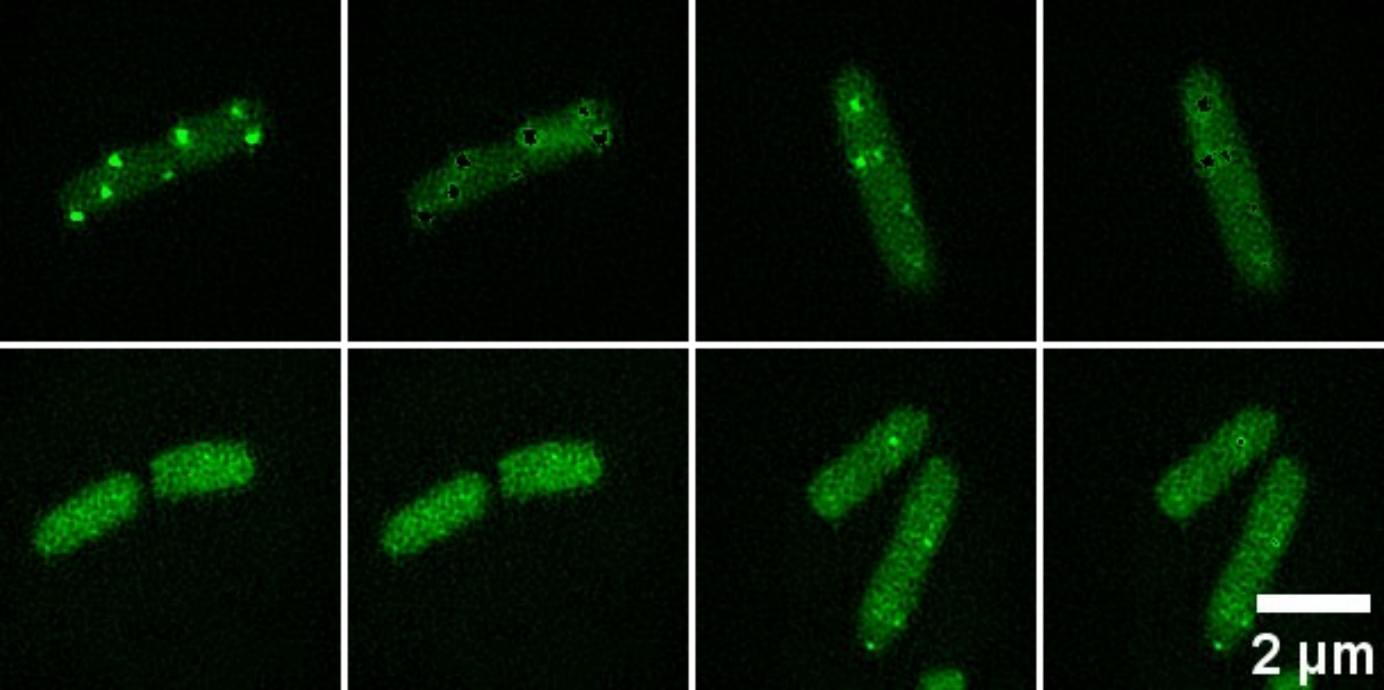

**Figure S2.** **Representative images of cells in the exponential phase (*OD*_600_ = 0.6) evaluated using a script developed for determining signal intensity percentage in fluorescent foci (Supplementary file 1).** Original images are shown in columns 1 and 3, while corresponding images in columns 2 and 4 were obtained after subtraction of the identified EcSSB foci.

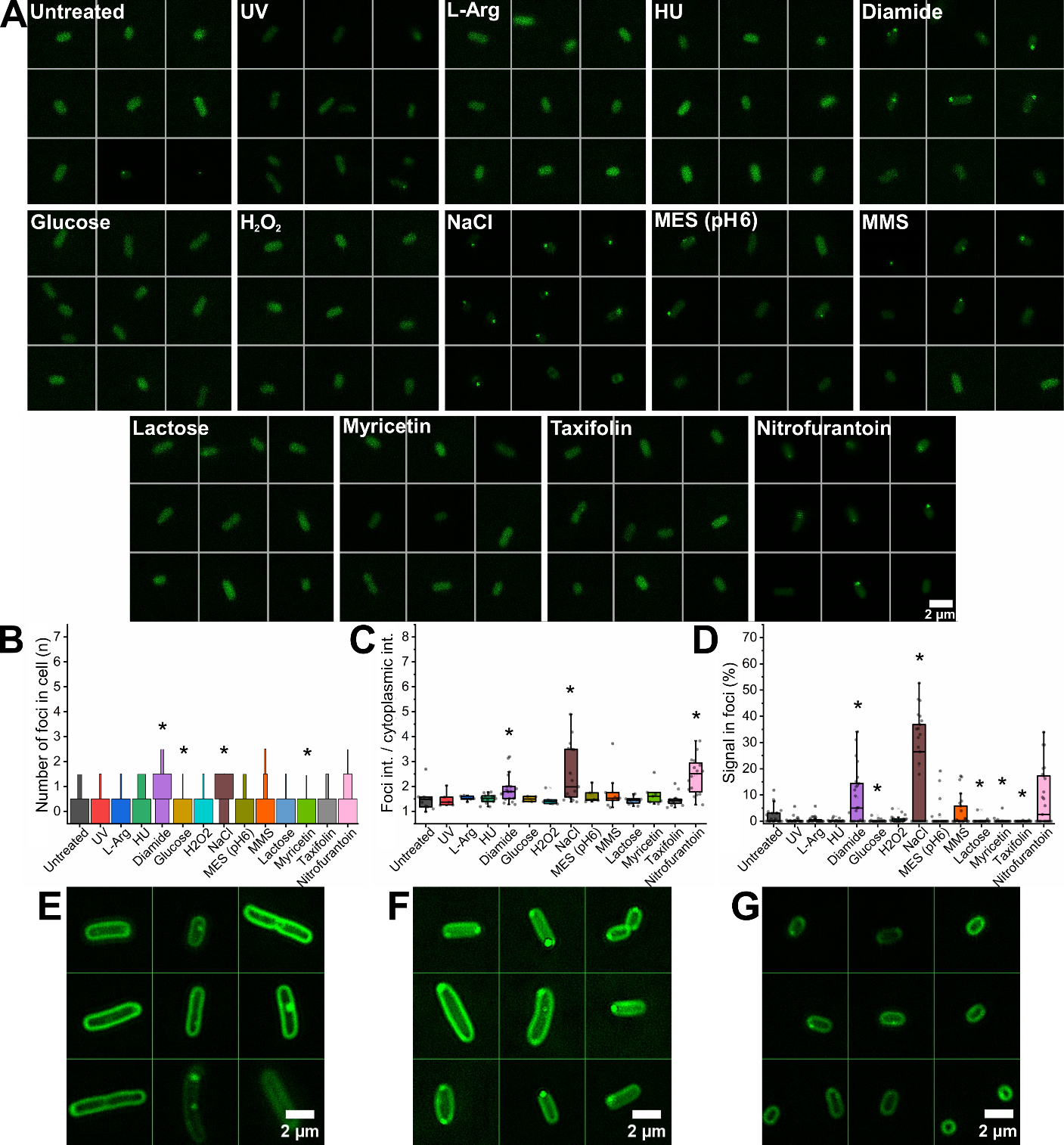

**Figure S3. Impact of various stressors on EcSSB-GFP foci during the death phase (OD600 = 0.6 + 24h). (A)** Representative SIM images of *E. coli* strain (#29003 ^1^) indicate that EcSSB-GFP foci rarely form during this phase. In addition, the overall EcSSB-GFP signal is weak, presumably due to reduced cellular EcSSB levels. Most stressors do not affect EcSSB organization patterns, with the exception of NaCl and diamide initiating foci formation. Stress induced by these agents could lead to local increase in EcSSB concentration, triggering assembly of EcSSB. **(B–D)** Only a few, less intense puncta (compared to those observed in earlier phases) are present, with EcSSB primarily localized homogeneously in the cytoplasm. 25 cells were analyzed per condition (see also **Table S5**). Statistical significance compared to untreated cells is denoted by black asterisks (Mann-Whitney test, *p* < 0.05). SIM images of cells in the **(E)** exponential, **(F)** stationary and **(G)** death phases treated with FM 4-64 membrane dye (8 µM).

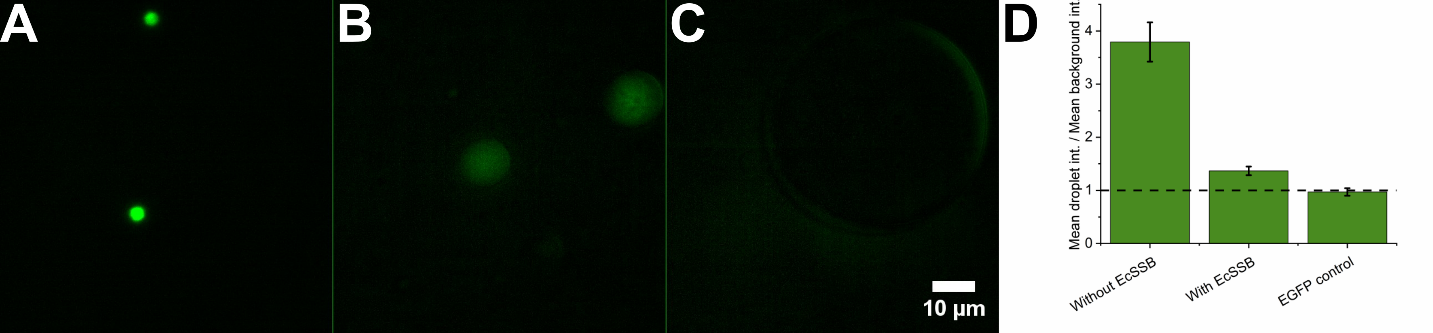

**Figure S4. EcSSB-GFP extracted from the #29003 cell line becomes readily embedded in WT EcSSB condensates.** (**A–B**) Fluorescence microscopy images of the supernatant from #29003 cells after lysis by sonication, diluted in buffer containing 20 mM Tris-acetate (pH 8.0), 50 mM KGlu, and 10 mM MgAc (**A**) without or (**B**) with the addition of 5 µM purified WT EcSSB tetramer. Note that the supernatant already contains spherical, bright droplets even without added EcSSB. Upon addition of EcSSB, droplets became larger and fainter. This behavior reflects spontaneous formation of EcSSB-GFP/WT EcSSB co-condensates and the free mixing of the two protein species in the droplets. (**C**) Control experiment showing EGFP signal in the same buffer containing 5 µM purified EcSSB tetramer without cell supernatant. (**D**) Intensity analysis of droplets and background showing accumulation of EcSSB-GFP, but not that of EGFP, in EcSSB condensates. Values greater than 1 indicate accumulation. Data represent mean ± SEM (*n* = 10).

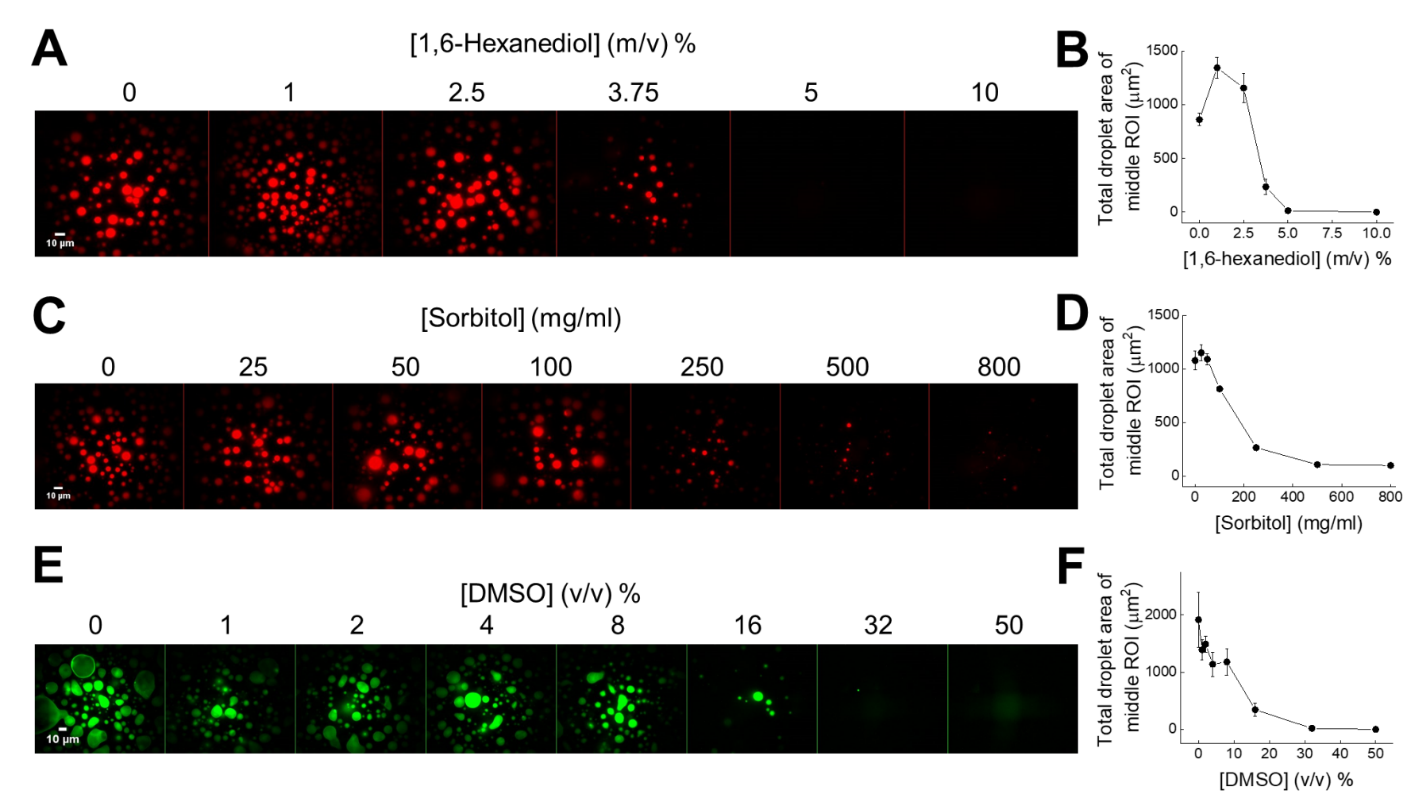

**Figure S5. *In vitro* response to chemical effectors corroborates liquid nature of EcSSB condensates. (A)** Representative epifluorescence microscopy images of EcSSB (9 µM, containing 0.15 µM AlexaFluor555-labeled EcSSB) mixed with indicated concentrations of 1,6-hexanediol, followed by 1-hour incubation. **(B)** Total droplet area of middle ROI from experiments shown in panel **A**. **(C)** Representative epifluorescence microscopy images of EcSSB (9 µM, containing 0.15 µM AlexaFluor555-labeled EcSSB) mixed with indicated concentrations of sorbitol, followed by 1-hour incubation. **(D)** Total droplet area of middle ROI from experiments shown in panel **C**. **(E)** Representative epifluorescence microscopy images of EcSSB (30 µM, containing 0.3 µM IAF-labeled EcSSB^G26C^) mixed with indicated concentrations of dimethyl sulfoxide (DMSO), followed by 1-min incubation. **(F)** Total droplet area of middle ROI from experiments shown in panel **E**. Error bars show SEM for *n* = 3 measurements on all panels.

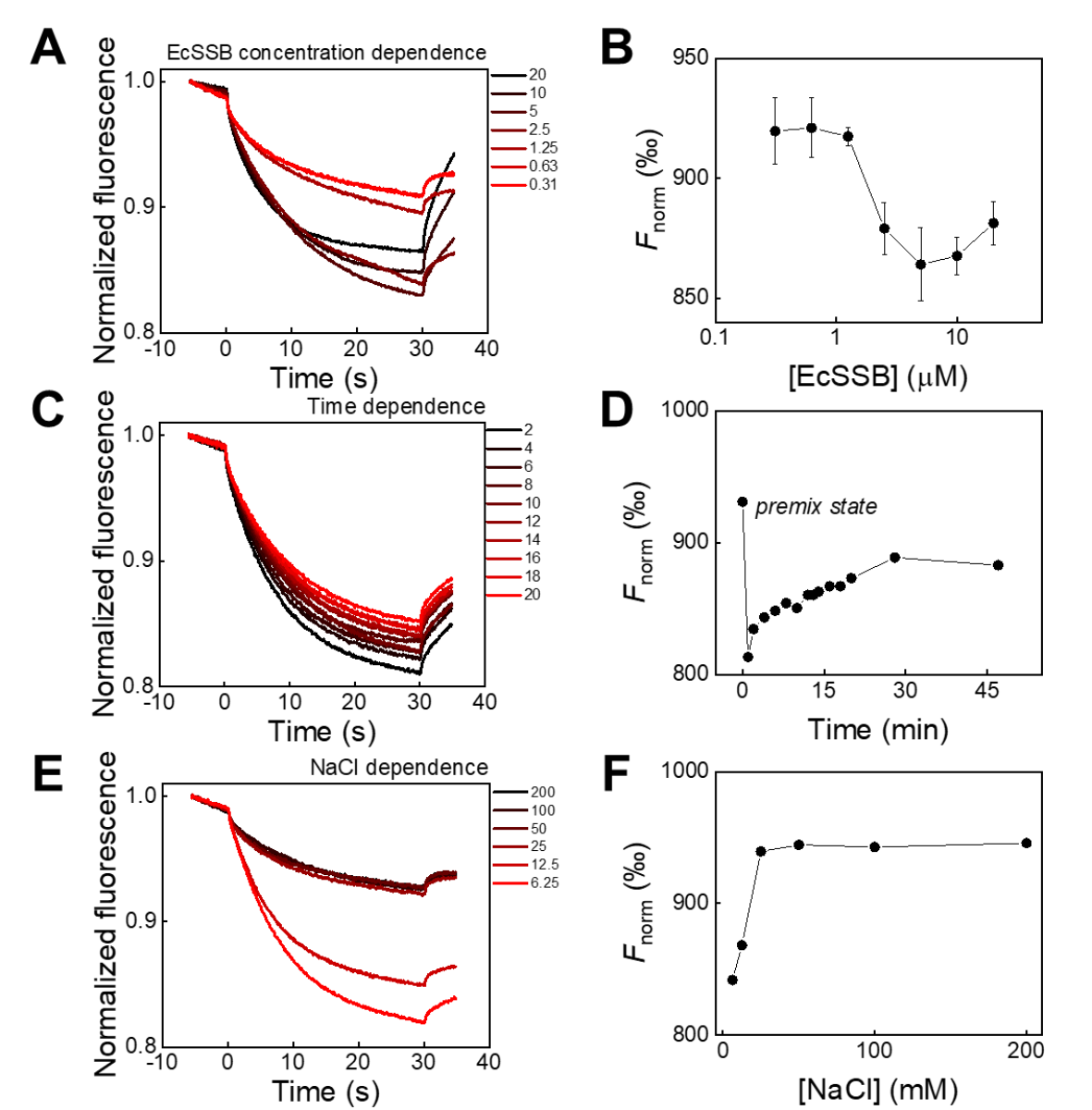

**Figure S6. Microscale thermophoresis (MST) is suitable for monitoring *in vitro* condensate formation, as thermophoretic mobility of EcSSB changes upon phase transition. (A)** MST traces of EcSSB samples of different EcSSB protein concentrations (indicated in µM; 50 nM IAF-labeled EcSSB^G26C^ was used in all cases). **(B)** Normalized fluorescence values (*F*_norm_), expressed as thousandths of the baseline fluorescence signal, measured at 20 seconds of irradiation in experiments shown in panel **A**. Error bars show SD of *n* = 2 independent measurements. **(C)** MST traces of 5 µM EcSSB samples (containing 50 nM IAF-labeled EcSSB^G26C^) incubated for increasing time periods (values indicated in minutes). **(D)** *F*_norm_ values (cf. panel **B**) from experiments shown in panel **C**. **(E)** NaCl concentration dependence (indicated in mM) of MST traces of 5 µM EcSSB (containing 50 nM IAF-labeled EcSSB^G26C^). **(F)** *F*_norm_ values (cf. panel **B**) from experiments shown in panel **E**.

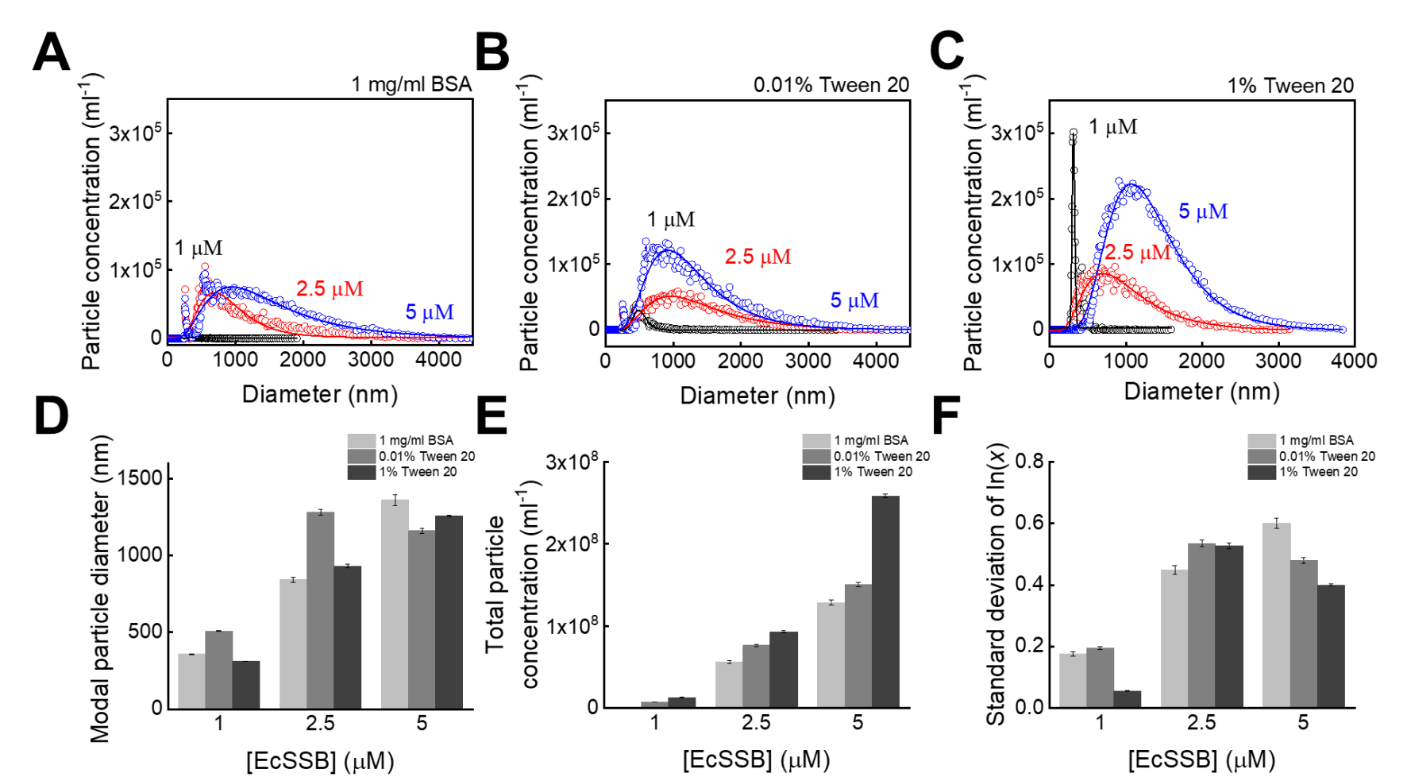

**Figure S7. Microfluidic resistive pulse sensing (MRPS) measurements reveal lognormal distribution of EcSSB condensate size. (A-C)** Size distribution of EcSSB condensates measured in LLPS buffer supplemented with **(A)** 1 mg/ml BSA, **(B)** 0.01 % Tween 20, and **(C)** 1 % Tween 20. Lines show best-fits with lognormal distribution. **(D)** Modal particle diameter, **(E)** total particle concentration and **(F)** standard deviation of the log-scale particle diameter values determined by lognormal fits shown in panels **A-C**. Error bars represent fitting standard errors. We note that cuvette blockage events were frequently encountered in conditions shown in panels **A-B**. Therefore, differences in particle size distributions and concentrations across different conditions **(D-F)** do not necessarily reflect the intrinsic features of EcSSB condensation. Cuvette blockage was negligible in 1 % Tween 20 (panel **C**). Therefore, this condition reflects most reliably the particle size distribution features of EcSSB **(D-F)**.

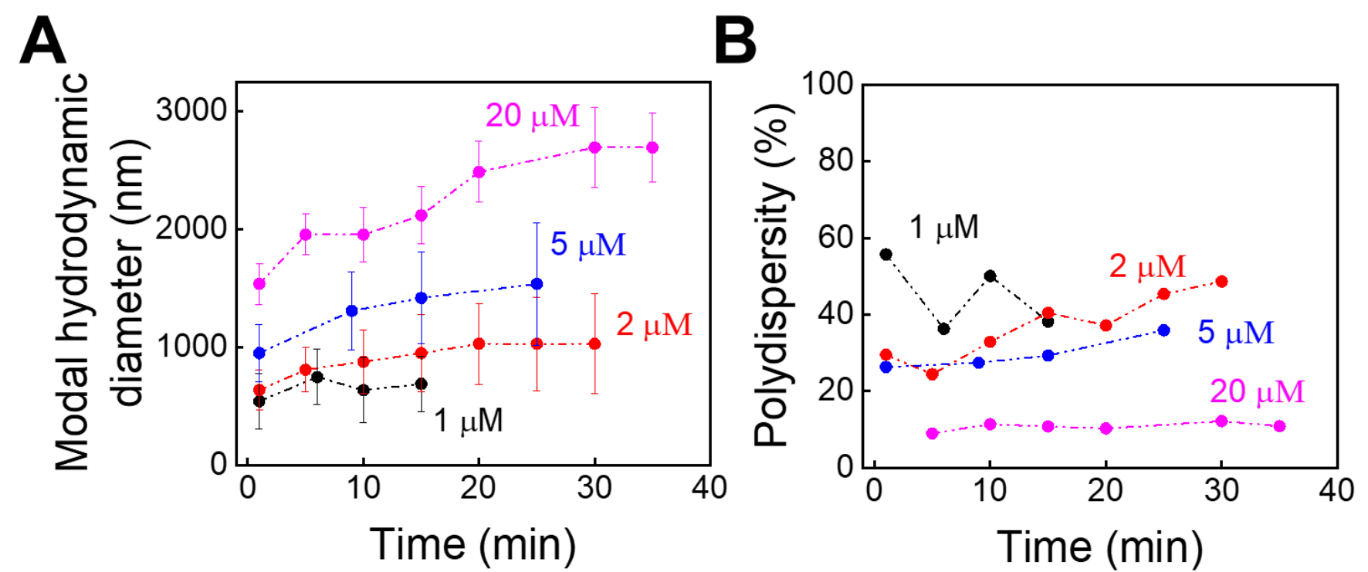

**Figure S8. Dynamic light scattering (DLS) experiments show time and protein concentration dependence of hydrodynamic diameter and polydispersity values of EcSSB condensates.** Time dependence is shown for modal hydrodynamic diameters (± SD) **(A)** and polydispersity **(B)** values of EcSSB samples (concentrations as indicated) in LLPS buffer measured *via* dynamic light scattering.

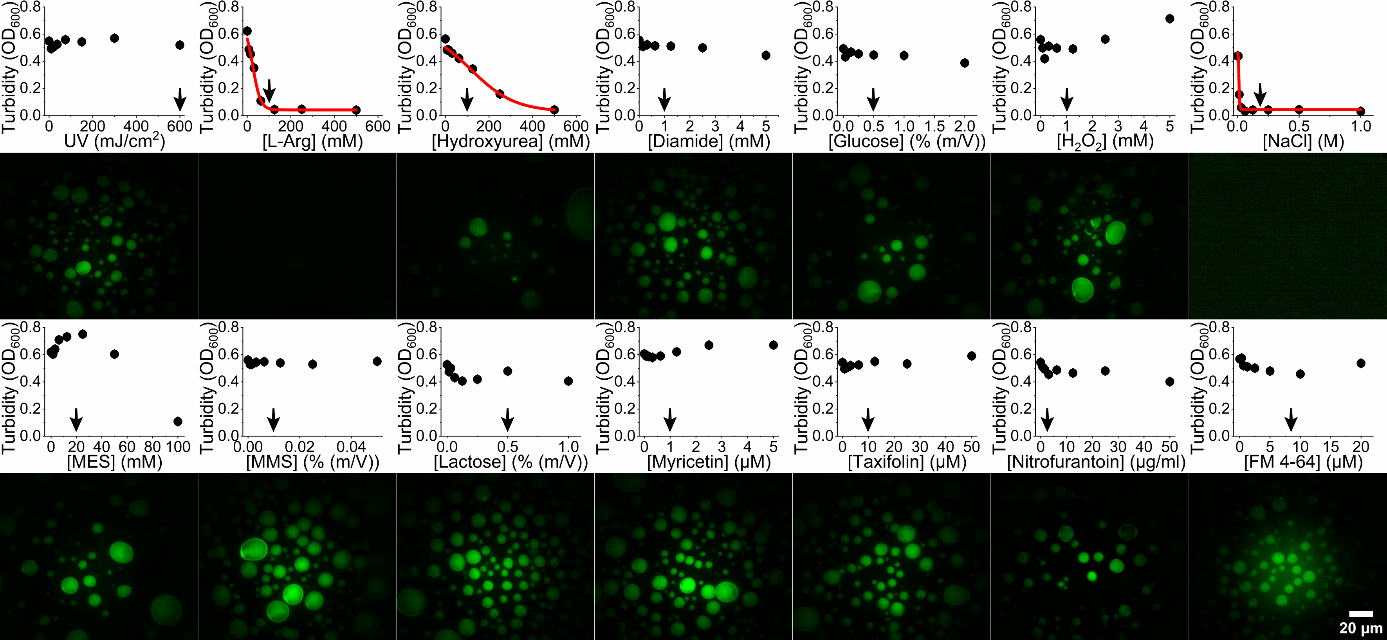

**Figure S9. *In vitro* turbidity measurements (top) and epifluorescence microscopy images (bottom) of EcSSB in the presence of stressors.** Turbidity signals decreasing with stressor concentrations are indicative of condensation inhibition ^2^. Black arrows indicate stressor concentrations applied in experiments *in vivo* (**Figs. 2-4, S2**; see also **Table S2**). Data points show mean ± SD of three independent measurements. A dose-response curve was fitted to data points in cases where stressors inhibited EcSSB condensate formation (*EC*_50_ ± SD: L-Arg, 25 ± 10 mM; HU, 120 ± 40 mM; NaCl, 13 ± 4 mM). Epifluorescence microscopy images show EcSSB condensates formed at stressor concentrations specified in **Table S2** (*in vitro* doses).

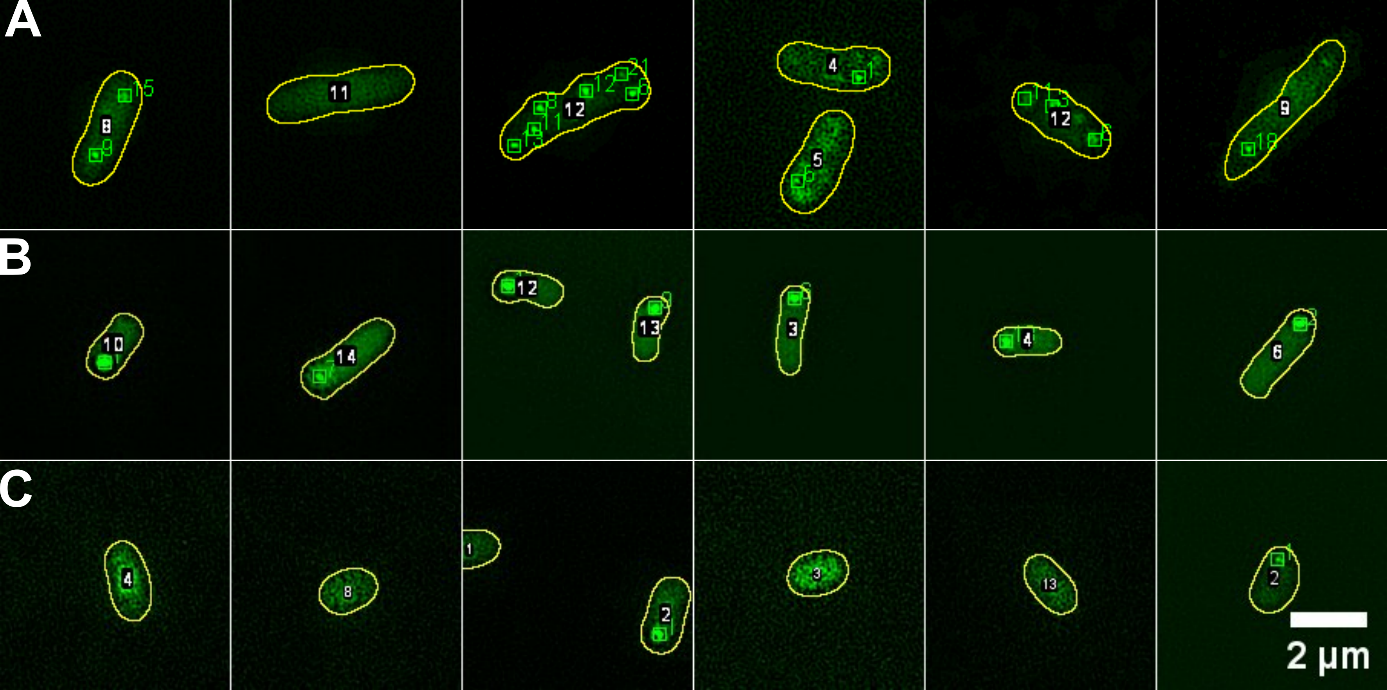

**Figure S10. Representation of the image evaluation method developed in this study.** Images were analyzed for cells in the **(A)** exponential (*OD*_600_ = 0.6), **(B)** stationary (*OD*_600_ = 0.6 + 3h) and **(C)** death (*OD*_600_ = 0.6 + 24h) phases. Cell outlines, as detected by the script, are marked in yellow, and the identified cells are labeled with white colored numbers. Foci identified within cells are marked as green squares and labeled with green colored numbers.

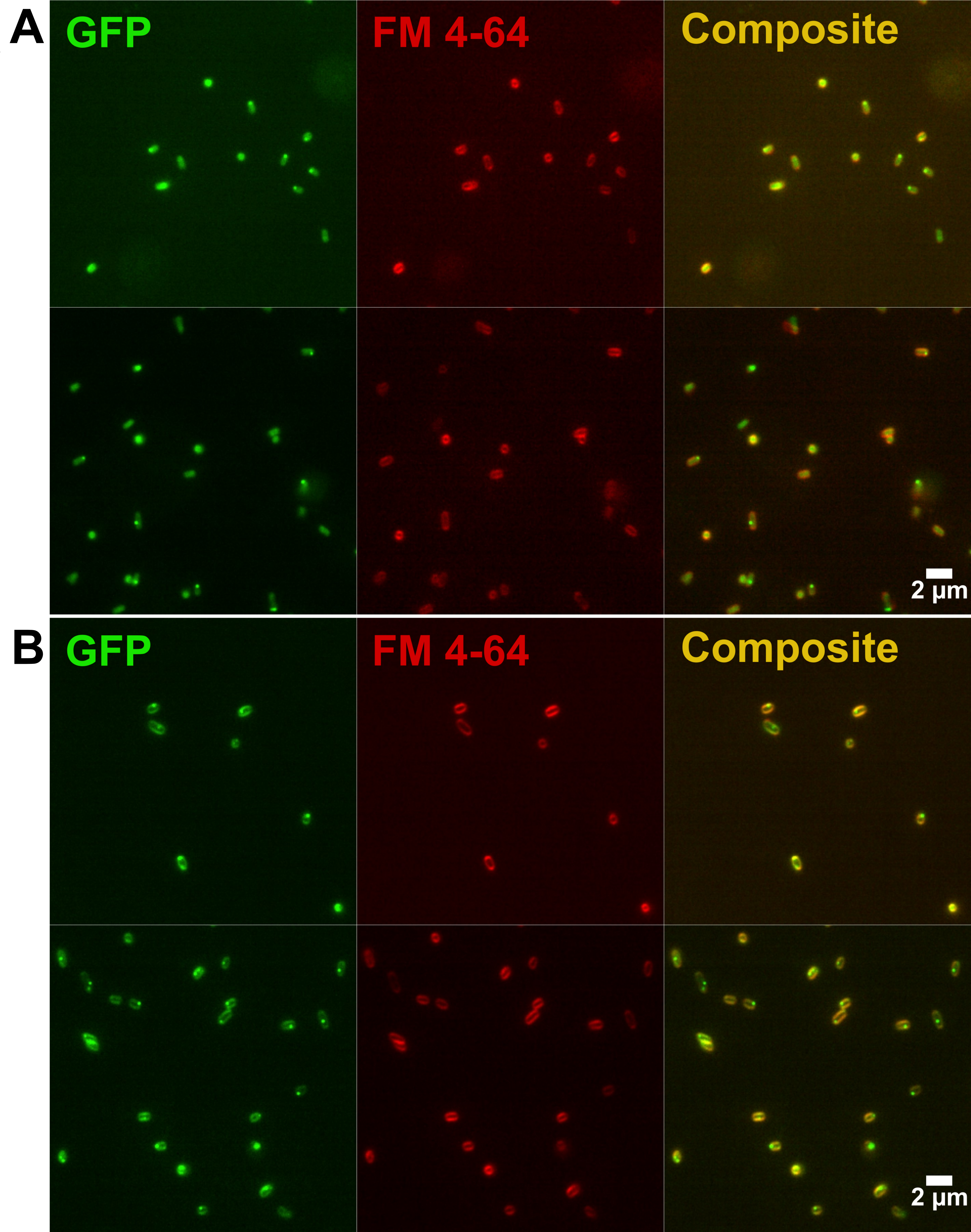

**Figure S11. FM 4-64 membrane dye promotes membrane localization of EcSSB-GFP at higher dye concentration.** Representative confocal images showing EcSSB-GFP (green) localization in the presence of FM 4-64 dye (red) applied at 2 µM (**A**) and 8 µM (**B**) concentrations.

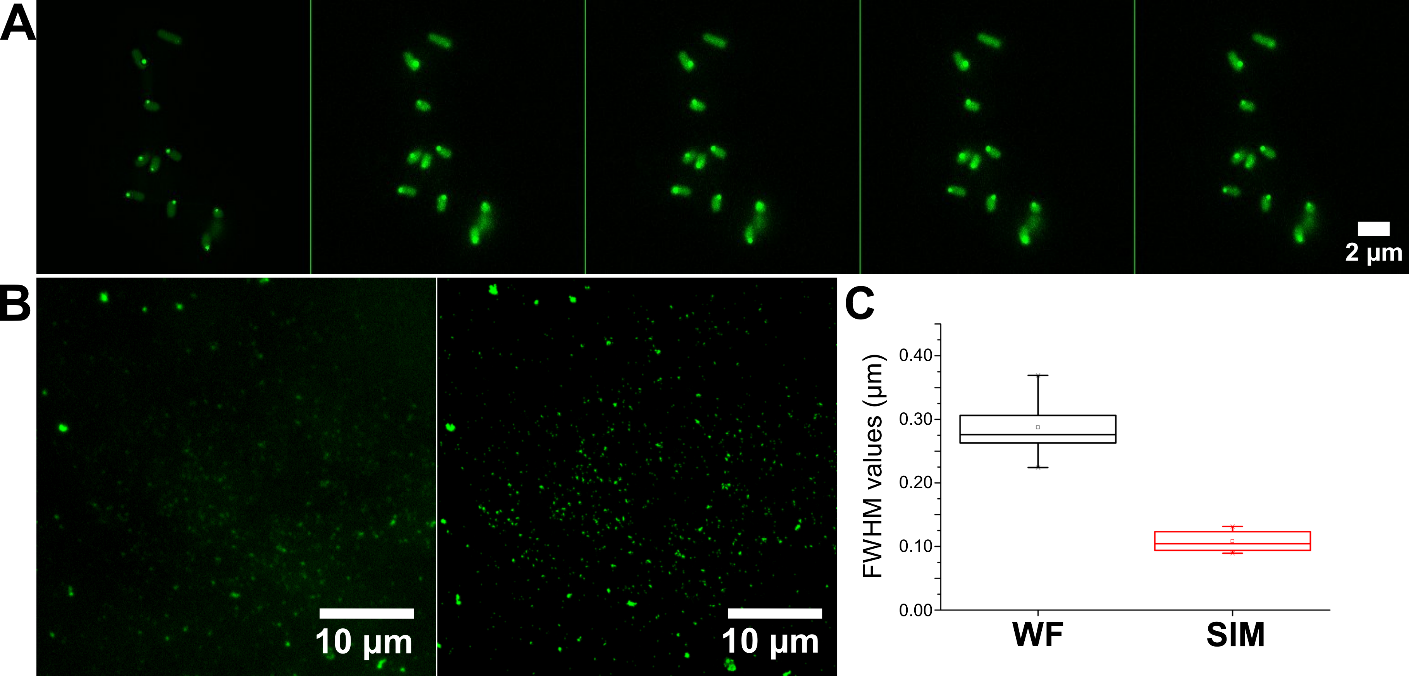

**Figure S12. Spatial resolution of SIM measurements.** (**A**) Comparison of processed (leftmost image) and unprocessed (other images to the right) SIM microscopy images of cells in the stationary phase. (**B**) Aberior Nanoparticles 4C fluor beads with an average diameter of 100 nm were used to determine the resolution of the microscope (left: wide-field (WF), right: SIM). (**C**) A Gaussian function were fitted to measured profiles of individual beads to determine their full width at half maximum (FWHM) values. WF values and those measured using the SIM method for the same beads differ significantly. In the case of SIM, the expected value of 100 nm was returned with a high degree of certainty.

**Table S1. Foci statistics in different growth phases (related to Fig. 1)**

| *OD*_600_ + time | Number of foci per cell | Foci int.  / cytoplasmic int. | | Signal in foci (%) | |
| --- | --- | --- | --- | --- | --- |
|  | Mode  (range, *n* of foci) | Median  (maximum) | Interquartile range (Q3 – Q1) | Median  (maximum) | Interquartile range (Q3 – Q1) |
| 0.2 | 0 (0-2, 25) | 1.41 (1.86) | 0.25 | 0 (15.35) | 0.73 |
| 0.6 | 1 (0-6, 95) | 2.00 (6.72) | 0.93 | 5.94 (33.27) | 9.34 |
| +1.5h | 1 (0-3, 62) | 2.38 (7.23) | 1.57 | 6.08 (56.53) | 19.53 |
| +3h | 1 (0-2, 66) | 3.88 (8.34) | 2.77 | 27.12 (64.70) | 35.47 |
| +4.5h | 1 (0-3, 57) | 5.13 (11.42) | 4.63 | 29.08 (63.55) | 38.81 |
| +7h | 1 (0-3, 43) | 6.28 (10.77) | 3.23 | 20.72 (66.27) | 32.54 |
| +10h | 0 (0-2, 32) | 5.97 (11.53) | 5.03 | 17.4 (69.35) | 37.05 |
| +24h | 0 (0-1, 14) | 1.47 (2.65) | 0.36 | 0 (12.85) | 0.23 |

Values are from 65 cells for each condition (*n* = 65).

**Table S2. Concentrations and physiological effects of stressors used in this study**

| Stressor (source) | *In vitro* dose | *In vivo* dose | Effects |
| --- | --- | --- | --- |
| UV  (UVP CX-2000 UV Crosslinker) | 600 mJ/cm^2^ | 600 mJ/cm^2^ | DNA damage: covalent pyrimidine base adducts; initiates SOS response ^3^^,^^4^ |
| L-Arg  (Sigma-Aldrich, A8094-25G) | 500 mM | 100 mM | Supports bacterial growth (nutrient) |
| Hydroxyurea  (Sigma-Aldrich, H8627-10G) | 500 mM | 100 mM | Reversibly inhibits ribonucleotide reductase and hence DNA synthesis ^5^ |
| Diamide  (Sigma-Aldrich, D3648-5G) | 5 mM | 1 mM | Thiol-specific thiol oxidizing agent, causes cysteine oxidation ^6^ |
| Glucose  (Sigma-Aldrich, G8270) | 2% | 0.5% | Supports bacterial growth (nutrient) |
| H_2_O_2_  (VWR Chemicals, 23619.297) | 5 mM | 1 mM | Oxidizing agent, DNA damage, protein denaturation ^7^ |
| NaCl  (VWR Chemicals, 27800.291) | 1 M | 200 mM | Osmotic stress; inhibits EcSSB phase separation *in vitro* ^2^ |
| MES (pH 6.0)  (Sigma-Aldrich, M2933-500G) | 100 mM | 20 mM | pH stress |
| MMS  (Sigma-Aldrich, 129925-5G) | 0.05% | 0.01% | Alkylating agent, induces DNA damage, affects replication, transcription ^8^ |
| Lactose  (Molar Chemicals Ltd.,  02150-203-190) | 2% | 0.5% | Supports bacterial growth (nutrient) |
| Myricetin  (Sigma-Aldrich, 72576-10MG) | 5 µM | 1 µM | Antimicrobial flavonoid ^9^; inhibits ssDNA binding activity of bacterial SSB ^10^ |
| Taxifolin  (Merck, 78666-25MG-F) | 50 µM | 10 µM | Antimicrobial flavonoid ^9^; inhibits ssDNA binding activity of bacterial SSB ^11^ |
| Nitrofurantoin  (Sigma-Aldrich, N7878-10G) | 50 µg/mL | 3 µg/mL | DNA damage: induces interstrand crosslinks ^12^ |
| FM 4-64  (Invitrogen, T13320) | 20 µM | 8 µM | Inner membrane dye ^1^ |

**Table S3. Effect of stressors on cells in the exponential phase (*OD*_600_ = 0.6)**

**(related to Fig. 2)**

|  | Number of foci per cell | | Foci int. / cytoplasmic int. | | | Signal in foci (%) | | | |
| --- | --- | --- | --- | --- | --- | --- | --- | --- | --- |
|  | Mode  (range,  *n* of foci) | Mann-Whitney *p* value | Median  (maximum) | Inter-quartile range  (Q3 – Q1) | Mann-Whitney *p* value | | Median  (maximum) | Inter-quartile range  (Q3 – Q1) | Mann-Whitney *p* value |
| Untreated | 1 (0-6, 88) |  | 2.00 (7.30) | 1.16 |  | | 4.06 (31.81) | 10.05 |  |
| UV | 0 (0-2, 38) | 2.2*10^-16^ | 1.73 (2.97) | 0.57 | 0.01 | | 0 (5.33) | 1.34 | 9.3*10^-8^ |
| L-Arg | 0 (0-2, 33) | 2.0*10^-7^ | 1.71 (3.30) | 0.58 | 0.03 | | 0 (17.64) | 2.68 | 8.3*10^-5^ |
| HU | 1 (0-5, 86) | 0.95 | 2.14 (6.15) | 1.13 | 0.82 | | 6.20 (39.26) | 13.07 | 0.28 |
| Diamide | 1 (0-4, 49) | 2.4*10^-6^ | 1.76 (3.50) | 0.47 | 7.1*10^-3^ | | 0 (21.26) | 4.13 | 3.2*10^-5^ |
| Glucose | 0 (0-3, 38) | 7.4*10^-10^ | 1.67 (3.18) | 0.49 | 0.02 | | 0.59 (9.70) | 2.08 | 3.5*10^-5^ |
| H_2_O_2_ | 1 (0-3, 52) | 1.1*10^-4^ | 1.73 (3.58) | 0.98 | 4.0*10^-3^ | | 0.41 (12.37) | 4.50 | 3.1*10^-4^ |
| NaCl | 1 (0-6, 126) | 0.02 | 2.38 (7.21) | 1.77 | 0.06 | | 9.37 (36.97) | 15.37 | 3.5*10^-3^ |
| MES (pH 6) | 0 (0-2, 19) | 1.0*10^-20^ | 1.71 (3.31) | 0.77 | 0.09 | | 0.31 (28.90) | 2.87 | 1.3*10^-3^ |
| MMS | 1 (0-4, 67) | 1.9*10^-3^ | 1.77 (3.65) | 0.38 | 9.8*10^-3^ | | 4.51 (30.78) | 11.55 | 0.78 |
| Lactose | 1 (0-3, 44) | 3.9*10^-6^ | 1.85 (3.83) | 1.05 | 0.03 | | 0.17 (23.68) | 3.95 | 4.9*10^-4^ |
| Myricetin | 0 (0-2, 34) | 2.0*10^-6^ | 1.58 (2.92) | 0.68 | 1.0*10^-3^ | | 0 (19.00) | 1.96 | 6.6*10^-6^ |
| Taxifolin | 0 (0-2, 38) | 1.9*10^-9^ | 1.61 (3.57) | 0.45 | 3.5*10^-4^ | | 0 (25.22) | 0.99 | 1.0*10^-6^ |
| Nitro-furantoin | 1 (0-5, 80) | 0.86 | 1.85 (7.09) | 1.28 | 0.68 | | 7.24 (35.67) | 14.67 | 0.61 |

Values are from 65 cells for each condition (*n* = 65).

**Table S4. Effect of stressors on cells in the stationary phase (*OD*_600_ = 0.6 + 3h)**

**(related to Fig. 3)**

|  | Number of foci per cell | | Foci int. / cytoplasmic int. | | | | Signal in foci (%) | | |
| --- | --- | --- | --- | --- | --- | --- | --- | --- | --- |
|  | Mode  (range, *n* of foci) | Mann-Whitney *p* value | Median  (maximum) | Inter-quartile range  (Q3 – Q1) | Mann-Whitney *p* value | Median  (maximum) | | Inter-quartile range  (Q3 – Q1) | Mann-Whitney *p* value |
| Untreated | 1 (0-2, 51) |  | 2.47 (7.62) | 2.38 |  | 20.43 (62.88) | | 21.80 |  |
| UV | 1 (0-5, 56) | 2.4*10^-9^ | 2.35 (5.60) | 1.39 | 0.09 | 6.10 (35.93) | | 15.83 | 4.2*10^-7^ |
| L-Arg | 1 (0-3, 73) | 9.9*10^-7^ | 2.84 (6.42) | 2.86 | 0.65 | 38.97 (54.69) | | 19.71 | 2.7*10^-5^ |
| HU | 1 (0-3, 40) | 0.03 | 2.11 (5.53) | 0.92 | 0.02 | 0 (47.25) | | 10.27 | 3.3*10^-10^ |
| Diamide | 1 (0-2, 59) | 0.05 | 2.93 (7.73) | 3.01 | 0.57 | 24.00 (63.55) | | 21.69 | 0.90 |
| Glucose | 1 (0-2, 51) | 0.83 | 2.80 (7.12) | 2.02 | 0.66 | 10.86 (48.07) | | 23.43 | 4.2*10^-3^ |
| H_2_O_2_ | 1 (0-4, 63) | 3.0*10^-4^ | 2.25 (4.90) | 1.27 | 0.03 | 8.74 (30.73) | | 16.01 | 1.0*10^-4^ |
| NaCl | 1 (0-2, 53) | 0.65 | 3.20 (8.77) | 2.39 | 0.35 | 15.76 (48.64) | | 30.29 | 0.05 |
| MES  (pH 6) | 1 (0-3, 74) | 4.8*10^-4^ | 2.56 (5.99) | 2.23 | 0.21 | 20.93 (48.79) | | 34.97 | 0.19 |
| MMS | 1 (0-3, 59) | 0.17 | 2.60 (6.62) | 2.16 | 0.38 | 14.19 (54.90) | | 28.49 | 0.10 |
| Lactose | 1 (0-3, 39) | 0.43 | 3.48 (7.25) | 2.82 | 0.23 | 11.00 (46.23) | | 22.85 | 3.3*10^-4^ |
| Myricetin | 1 (0-2, 64) | 0.11 | 3.02 (8.20) | 1.84 | 0.41 | 20.95 (52.48) | | 29.61 | 0.42 |
| Taxifolin | 1 (0-2, 61) | 0.20 | 2.62 (10.58) | 1.93 | 0.51 | 15.17 (52.50) | | 28.94 | 0.10 |
| Nitro-furantoin | 1 (0-2, 57) | 0.30 | 3.22 (6.97) | 3.35 | 0.31 | 19.00 (57.77) | | 37.36 | 0.49 |

Values are from 65 cells for each condition (*n* = 65).

**Table S5. Effect of stressors on cells in the death phase (*OD*_600_ = 0.6 + 24h)**

**(related to Fig. S2)**

|  | Number of foci per cell | | Foci int. / cytoplasmic int. | | | | Signal in foci (%) | | |
| --- | --- | --- | --- | --- | --- | --- | --- | --- | --- |
|  | Mode  (range, *n* of foci) | Mann-Whitney *p* value | Median  (maximum) | Inter-quartile range  (Q3 – Q1) | Mann-Whitney *p* value | Median  (maximum) | | Inter-quartile range  (Q3 – Q1) | Mann-Whitney *p* value |
| Untreated | 0 (0-1, 11) |  | 1.48 (2.69) | 0.39 |  | 0 (11.81) | | 3.07 |  |
| UV | 0 (0-2, 5) | 0.25 | 1.37 (2.03) | 0.31 | 0.73 | 0 (5.63) | | 0.24 | 0.06 |
| L-Arg | 0 (0-1, 5) | 0.17 | 1.51 (1.64) | 0.12 | 0.31 | 0 (5.70) | | 0.55 | 0.12 |
| HU | 0 (0-1, 14) | 0.26 | 1.52 (1.77) | 0.22 | 0.21 | 0 (1.82) | | 0.23 | 0.06 |
| Diamide | 1 (0-4, 18) | 5.3*10^-6^ | 1.79 (3.19) | 0.49 | 0.01 | 5.02 (34.10) | | 14.38 | 0.02 |
| Glucose | 0 (0-1, 4) | 0.01 | 1.49 (1.60) | 0.20 | 0.32 | 0 (2.24) | | 0 | 0.01 |
| H_2_O_2_ | 0 (0-1, 7) | 0.12 | 1.39 (1.94) | 0.15 | 1 | 0 (4.77) | | 0.90 | 0.47 |
| NaCl | 1 (0-1, 17) | 4.2*10^-6^ | 1.98 (4.89) | 1.91 | 4.1*10^-3^ | 26.43 (52.61) | | 36.85 | 9.5*10^-3^ |
| MES  (pH 6) | 0 (0-1, 5) | 0.56 | 1.47 (2.15) | 0.31 | 0.42 | 0 (19.14) | | 0 | 0.08 |
| MMS | 0 (0-2, 9) | 0.55 | 1.52 (3.71) | 0.31 | 0.28 | 0 (17.18) | | 5.80 | 0.73 |
| Lactose | 0 (0-1, 11) | 0.09 | 1.44 (1.70) | 0.17 | 0.94 | 0 (4.33) | | 0 | 4.0*10^-3^ |
| Myricetin | 0 (0-1, 7) | 0.02 | 1.61 (2.56) | 0.38 | 0.17 | 0 (5.01) | | 0 | 5.1*10^-3^ |
| Taxifolin | 0 (0-1, 13) | 0.50 | 1.42 (2.11) | 0.16 | 1 | 0 (0.44) | | 0 | 9.8*10^-4^ |
| Nitro-furantoin | 0 (0-2, 16) | 0.11 | 2.51 (3.82) | 1.1 | 4.1*10^-4^ | 2.58 (33.92) | | 17.22 | 0.11 |

Values are from 25 cells for each condition (*n* = 25).

**SUPPLEMENTARY FILE 1**

**Protocol for Foci Analysis**

1. Place microscopy images of interest in a designated folder.
2. In the Fiji macro window, load the file named *Fiji_macro_for_foci_analyses.ijm.* Specify the folder path where images are located.
3. Execute the macro.
4. Check the output files:

For each microscopy image the macro will generate a .txt file, a .tif file, and two Excel files. These will share the same name as the original image file.

The .tif file shows regions identified as ROI and foci classified. Use these images to fine-tune the macro for more accurate foci detection (lines 62–63 and 184–185 in the script) and ROI detection by optimizing thresholds (line 144 in the macro).

5. Use the following RStudio scripts to merge the generated Excel and .txt files:
   - *Cell_area_and_average_intensity*
   - *Number_of_foci_in_cells*
   - *Position_and_intensity_of_foci*

Merged files will display data for each image in separate sections, labeled accordingly.

**Protocol for Subtracting Foci from Intensity Measurement**

1. Execute the *Fiji_macro_for_foci_percentage.ijm* macro in Fiji.
2. Check the output files:

For each image file, two Excel files will be generated. These contain the average intensity values for each cell with foci and without foci.

Note: Thresholds must be optimized to suit the microscopy images (lines 45 and 63 in the script).

3. Use the following RStudio scripts to merge the data:
   - *Cell_intensity_with_foci*
   - *Cell_intensity_without_foci*

Merged files will provide average intensity values for cells with and without foci.

**References**

1	Zhao, T. *et al.* Super-resolution imaging reveals changes in Escherichia coli SSB localization in response to DNA damage. *Genes to cells : devoted to molecular & cellular mechanisms* **24**, 814-826, doi:10.1111/gtc.12729 (2019).

2	Harami, G. M. *et al.* Phase separation by ssDNA binding protein controlled via protein-protein and protein-DNA interactions. *Proceedings of the National Academy of Sciences of the United States of America* **117**, 26206-26217, doi:10.1073/pnas.2000761117 (2020).

3	Harami, G. M. *et al.* Shuttling along DNA and directed processing of D-loops by RecQ helicase support quality control of homologous recombination. *Proceedings of the National Academy of Sciences of the United States of America* **114**, E466-E475, doi:10.1073/pnas.1615439114 (2017).

4	Yang, W. Surviving the sun: repair and bypass of DNA UV lesions. *Protein science : a publication of the Protein Society* **20**, 1781-1789, doi:10.1002/pro.723 (2011).

5	Singh, A. & Xu, Y. J. The Cell Killing Mechanisms of Hydroxyurea. *Genes* **7**, doi:10.3390/genes7110099 (2016).

6	Kosower, N. S. & Kosower, E. M. Diamide: an oxidant probe for thiols. *Methods in enzymology* **251**, 123-133, doi:10.1016/0076-6879(95)51116-4 (1995).

7	Juven, B. J. & Pierson, M. D. Antibacterial Effects of Hydrogen Peroxide and Methods for Its Detection and Quantitation (dagger). *Journal of food protection* **59**, 1233-1241, doi:10.4315/0362-028X-59.11.1233 (1996).

8	Volkert, M. R. & Landini, P. Transcriptional responses to DNA damage. *Current opinion in microbiology* **4**, 178-185, doi:10.1016/s1369-5274(00)00186-7 (2001).

9	Shamsudin, N. F. *et al.* Antibacterial Effects of Flavonoids and Their Structure-Activity Relationship Study: A Comparative Interpretation. *Molecules* **27**, doi:10.3390/molecules27041149 (2022).

10	Lin, E. S., Luo, R. H. & Huang, C. Y. A Complexed Crystal Structure of a Single-Stranded DNA-Binding Protein with Quercetin and the Structural Basis of Flavonol Inhibition Specificity. *International journal of molecular sciences* **23**, doi:10.3390/ijms23020588 (2022).

11	Lin, E. S., Huang, Y. H., Luo, R. H., Basharat, Z. & Huang, C. Y. Crystal Structure of an SSB Protein from Salmonella enterica and Its Inhibition by Flavanonol Taxifolin. *International journal of molecular sciences* **23**, doi:10.3390/ijms23084399 (2022).

12	Sengupta, S. *et al.* DNA damage and prophage induction and toxicity of nitrofurantoin in Escherichia coli and Vibrio cholerae cells. *Mutation research* **244**, 55-60, doi:10.1016/0165-7992(90)90108-v (1990).
