## Supplementary material for "Phase transition of bacterial single-stranded DNA binding (SSB) protein upon stress response and metabolic adaptation": Scripts: Cell_area_and_average_intensity.R

```
# Installing and loading necessary packages
install.packages("writexl")
library(writexl)

# Path to the folder where the CSV files are located
folder <- "F:/"

# Create a list of CSV files that end with "spots"
csv_files <- list.files(path = folder, pattern = ".czi\\.csv$", full.names = TRUE)

# Check if any files were found
if (length(csv_files) == 0) {
  stop("No CSV files ending with 'SD.nd2' found in the specified folder.")
}

# Create an empty data frame
data_table <- data.frame()

# Loop through the CSV files
for (file in csv_files) {
  # Determine the name of the CSV file
  file_name <- tools::file_path_sans_ext(basename(file))
  
  # Read the CSV file
  data <- read.csv(file, stringsAsFactors = FALSE)
  
  # Add the data to the data table
  data_table <- rbind(data_table, cbind(file_name, data))
}

# Create an Excel file and save the data table
write_xlsx(data_table, file.path(folder, "Cell_area_and_average_intensity.xlsx"))
```
