## Supplementary material for "Phase transition of bacterial single-stranded DNA binding (SSB) protein upon stress response and metabolic adaptation": Scripts: Fiji_macro_for_foci_analyses.ijm

```
requires("1.33s");
//Set the folder where the images can be found
   dir = getDirectory("F:/");
   setBatchMode(true);
   count = 0;
   countFiles(dir);
   n = 0;
   processFiles(dir);
   //print(count+" files processed");

   function countFiles(dir) {
      list = getFileList(dir);
      for (i=0; i<list.length; i++) {
          if (endsWith(list[i], "/"))
              countFiles(""+dir+list[i]);
          else
              count++;
      }
  }

   function processFiles(dir) {
      list = getFileList(dir);
      for (i=0; i<list.length; i++) {
          if (endsWith(list[i], "/"))
              processFiles(""+dir+list[i]);
          else {
             showProgress(n++, count);
             path = dir+list[i];
             processFile(path);
          }
      }
  }

  function processFile(path) {
       if (endsWith(path, ".czi")) {
           open(path);

run("Duplicate...", "title=ImageTitle");
selectWindow("ImageTitle");

//Find foci and show them

run("Duplicate...", "title=original duplicate channels=1-2");
selectWindow("original");

run("Duplicate...", "title=selected duplicate channels=1-2");
selectWindow("selected");

run("Duplicate...", "title=background duplicate channels=1-2");
selectWindow("original");
selectWindow("background");

run("Set Measurements...", "standard modal"); 
List.setMeasurements(); 
ch1modevalue=List.getValue("Mode");
ch1sdvalue=List.getValue("StdDev"); 

run("Max...", "value="+ch1modevalue);
run("Gaussian Blur...", "sigma=100");

// These values can and should be optimized depending on the intensity of foci
ch1selval=ch1sdvalue*22;
radius=4;

run("Line Width...", "line=1");

selectWindow("original");
setSlice(1); 

run("Find Maxima...", "noise="+ch1selval+" output=List exclude");

run("ROI Manager...");	
roiManager("reset"); 
run("Set Measurements...", "  mean redirect=None decimal=1");

String.resetBuffer;
String.copyResults;
filestring=String.paste;

rows=split(filestring, "\n");
particle=newArray(rows.length); 
x=newArray(rows.length); 
y=newArray(rows.length); 

print("prtcl","\t","\t","xpos","\t","\t","ypos","\t","\t","ch1val");

for(i=0; i<rows.length; i++){ 
columns=split(rows[i],"\t"); 
particle[i]=parseInt(columns[0]);
x[i]=parseInt(columns[1]); 
y[i]=parseInt(columns[2]); 
makeRectangle(x[i]-radius, y[i]-radius, radius*2, radius*2);
setLineWidth(10);
setForegroundColor(255, 255, 255);
roiManager("Add");
roiManager("Multi Measure");
ch1val=getResult("Mean1", 0);

print (particle[i],"\t","\t",x[i],"\t","\t",y[i],"\t","\t",ch1val);

	logforsave = path+"spotok.txt";
	selectWindow("Log");
	saveAs("Text", logforsave);
	roiManager("reset");
}

// Draw in the selected particles
selectWindow("selected");

for(i=0; i<rows.length; i++){ 
columns=split(rows[i],"\t"); 
particle[i]=parseInt(columns[0]);
x[i]=parseInt(columns[1]); 
y[i]=parseInt(columns[2]); 
radius=4;
makeRectangle(x[i]-radius, y[i]-radius, radius*2, radius*2);
setLineWidth(300);
setForegroundColor(255, 255, 255);
setFont("Arial", 15);
setForegroundColor(255, 255, 255);
drawString(particle[i],x[i]+radius,y[i]+radius);
run("Draw");
} 

// Collecting Log results
selectWindow("Log");
print("\\Clear");

//setBatchMode(false);


//Find cells
	selectImage("ImageTitle");
	
	//run("Set Measurements...", "area mean integrated median display redirect="+ImageTitle+" decimal=2");
	
	run("Duplicate...", " ");
	setMinAndMax(0, 30000);
	run("Apply LUT");
	run("Subtract Background...", "rolling=50");
	run("Gaussian Blur...", "sigma=5");
	run("Gamma...", "value=1.5");
//Tresholding should be optimized to find the whole cell
	setAutoThreshold("Li dark");
	run("Convert to Mask");
	run("Despeckle");
	run("Fill Holes");
	run("Open");
	run("Close-");
	run("Set Measurements...", "area mean standard redirect=None decimal=1");
	run("Analyze Particles...", "size=100-Infinity pixel add");
	close();
	roiManager("Show all");
	roiManager("Measure");
	resultsforsave = path+".csv";
	saveAs("Results", resultsforsave);
		
//Thresholding for LLPS inside the cell
		run("Duplicate...", " ");
		run("Set Measurements...", "standard"); 
		List.setMeasurements(); 
		ch1sdvalue=List.getValue("StdDev");
		run("Max...", "value="+ch1modevalue);
		close();
		
		run("Duplicate...", " ");
		selectWindow("ImageTitle");
		run("Duplicate...", "title=original2");
		selectWindow("original2");
		run("Duplicate...", "title=background2");
		selectWindow("original2");
		selectWindow("background2");
		
		run("Set Measurements...", "standard"); 
		List.setMeasurements(); 

		run("Max...", "value="+ch1modevalue);
		run("Gaussian Blur...", "sigma=100");
		run("Line Width...", "line=1");
		
		ch1sdvalue=List.getValue("StdDev");
		run("Clear Results");
// These values can and should be otimized depending on the intensity of foci (also above) 		 
		ch1selval=ch1sdvalue*22;
		radius=4;

		selectWindow("ImageTitle");
{
c= roiManager("count");
for (i = 0; i < c; i++) {
		roiManager("Select", i);
		run("Find Maxima...", "noise="+ch1selval+" output=Count");		
 	}
 }
	log_count_forsave = path+"counted_spots.csv";
	saveAs("Results", log_count_forsave);
	
selectImage("background");
close();
selectImage("original");
close();
selectWindow("original2");
close();
selectWindow("background2");
close();
selectWindow("ImageTitle-1");
close();
selectWindow("ImageTitle");
close();
selectImage("selected");
roiManager("Show all");
run("Flatten");

pathforsave = path+".tif";
saveAs("Tiff", pathforsave);

roiManager("Deselect");
roiManager("Delete");
run("Close All");
	}
 }
```
