## Supplementary material for "Phase transition of bacterial single-stranded DNA binding (SSB) protein upon stress response and metabolic adaptation": Scripts: Fiji_macro_for_foci_percentage.ijm

```
//Set the folder where the images can be found
dir = getDirectory("F:/");
setBatchMode(true);
count = 0;
countFiles(dir);
n = 0;
processFiles(dir);
//print(count+" files processed");

function countFiles(dir) {
   list = getFileList(dir);
   for (i = 0; i < list.length; i++) {
       if (endsWith(list[i], "/"))
           countFiles("" + dir + list[i]);
       else
           count++;
   }
}

function processFiles(dir) {
   list = getFileList(dir);
   for (i = 0; i < list.length; i++) {
       if (endsWith(list[i], "/"))
           processFiles("" + dir + list[i]);
       else {
          showProgress(n++, count);
          path = dir + list[i];
          processFile(path);
       }
   }
}

function processFile(path) {
    if (endsWith(path, ".czi")) {
        open(path);
//Detecting the whole cell (optimization might be needed)        
        rename("Original_Image");
        run("Duplicate...", "title=Processed_Image");
        setMinAndMax(0, 30000);
        run("Apply LUT");
        run("Subtract Background...", "rolling=50");
        run("Gaussian Blur...", "sigma=5");
        run("Gamma...", "value=1.5");
//Tresholding should be optimized to detect the whole cell
        setAutoThreshold("Triangle dark");
        run("Convert to Mask");
        run("Despeckle");
        run("Fill Holes");
        run("Open");
        run("Close-");
        run("Set Measurements...", "area mean standard integrated redirect=None decimal=1");
        run("Analyze Particles...", "size=100-Infinity pixel add");
        close();
        roiManager("Show all");
        roiManager("Measure");
        resultsforsave = path + "withfoci.csv";
        saveAs("Results", resultsforsave);
        run("Clear Results");
        close("Results");

// Applying threshold to detect foci (treshold should be optimized)
        run("Duplicate...", "title=Moments_Threshold");
        setAutoThreshold("Yen dark");

// Exclude areas outside the threshold (invert and apply selection)
        run("Create Selection");
// Reset areas outside the selection        
        run("Clear Outside");
        selectImage("Original_Image");
        run("Duplicate...", "title=Processed_Image2");
        run("Image Calculator...", "image1=Processed_Image2 operation=Subtract image2=Moments_Threshold");

        selectImage("Original_Image");
        close();
        selectImage("Moments_Threshold");
        close();
        selectImage("Processed_Image2");
        rename("Original_Image");

        roiManager("Show all");
        roiManager("Measure");
        resultsforsave = path + "withoutfoci.csv";
        saveAs("Results", resultsforsave);
        close("Results");

        roiManager("Deselect");
        roiManager("Delete");
        run("Close All");
    }
}
```
