## Supplementary material for "Phase transition of bacterial single-stranded DNA binding (SSB) protein upon stress response and metabolic adaptation": Scripts: Number_of_foci_in_cells.R

```
# Installing required packages
install.packages("writexl")
library(writexl)

# Directory path where the CSV files are located
mappa <- "F:/"


csv_files <- list.files(path = mappa, pattern = "spots\\.csv$", full.names = TRUE)

# Checking if any matching files are found
if (length(csv_files) == 0) {
  stop("No CSV files ending with 'spots' were found in the specified directory.")
}

# Creating an empty data table
adattabla <- data.frame()

# Iterating through the CSV files
for (file in csv_files) {
  # Determining the name of the CSV file
  file_nev <- tools::file_path_sans_ext(basename(file))
  
  # Reading the CSV file
  adatok <- read.csv(file, stringsAsFactors = FALSE)
  
  # Adding data to the data table
  adattabla <- rbind(adattabla, cbind(file_nev, adatok))
}

# Creating an Excel file and saving the data table
write_xlsx(adattabla, file.path(mappa, "Number_of_foci_in_cells.xlsx"))
```
