## Supplementary material for "Phase transition of bacterial single-stranded DNA binding (SSB) protein upon stress response and metabolic adaptation": Scripts: Position_and_intensity_of_foci.R

```
# Installing required packages
install.packages("writexl")
library(writexl)

# Directory path where the TXT files are located
directory <- "F:/"

# Creating a list of TXT files
txt_files <- list.files(path = directory, pattern = "\\.txt$", full.names = TRUE)

# Creating an empty data table
data_table <- data.frame()

# Iterating through the TXT files
for (file in txt_files) {
  # Determining the name of the TXT file
  file_name <- tools::file_path_sans_ext(basename(file))
  
  # Reading the TXT file line by line
  lines <- readLines(file, warn = FALSE)
  
  # Splitting each line into columns using a delimiter (e.g., tab or comma)
  # Here, assuming a tab delimiter. Change `"\t"` to `","` for comma-separated values.
  split_lines <- do.call(rbind, strsplit(lines, "\t"))
  
  # Converting the split lines to a data frame
  data_frame <- data.frame(file_name, split_lines, stringsAsFactors = FALSE)
  
  # Adding data to the main data table
  data_table <- rbind(data_table, data_frame)
}

# Creating an Excel file and saving the data table
write_xlsx(data_table, file.path(directory, "Position_and_intensity_of_foci.xlsx"))
```
